# Modulation of miR-23b Wnt/β-catenin Axis Strengthens Endothelial Barrier Properties

**DOI:** 10.1101/2025.08.20.671398

**Authors:** Victor Anthony Martinez, Rose Presby, Sara Nakanishi, Pallavi Gaur, Ugur Akcan, Johannes Daugaard Krog, Iben Daugaard Krog, Lipi Das, David S. Johnson, Danny Jamoul, Douglas Jury, Brian Lawson, Jacob Körbelin, Lewis M. Brown, Vilas Menon, Dritan Agalliu, Irene Munk Pedersen

## Abstract

Blood-brain barrier (BBB) disruption drives stroke and other CNS disorders pathology, yet restoring its integrity remains challenging. Using an unbiased anti-miR lentiviral screen, miR-23b was identified as a critical negative regulator of BBB integrity in brain endothelial cells (BECs). Targeted inhibition of miR-23b using anti-miR-23b, validated by Wnt-reporter, Wnt-rescue, and eCLIP analysis, demonstrates that BEC miR-23b silencing in BECs, enhances junctional protein expression and strengthens barrier function while suppressing transcellular transport. Global multi-omics analysis reveals that miR-23b acts as a robust driver of angiogenesis-specific genes and proteins. Conversely, anti-miR-23b induces the expression of factors essential for BBB stabilization and repair. Notably, anti-miR-23b enhances these protective barrier properties through the multi-faceted regulation of Wnt/β-catenin, TGF-β, Notch, and VEGF signaling mechanisms. Using a 3D microfluidic platform, anti-miR-23b was shown to foster BBB repair by accelerating vessel maturation and enhancing resilience against ischemic injury. Proof-of-concept studies show that BEC specific AAVBR1-anti-miR-23b gene therapy reinforces BBB tight junctions, reduces BBB leakage, and improves outcome measures in a transient middle cerebral artery occlusion (t-MCAO) stroke model. Thus, miR-23b is a critical regulator of cerebrovascular integrity, positioning anti-miR-23b as a promising RNA-based therapy to enhance BBB stability and repair in stroke and other CNS disorders.

## INTRODUCTION

Neurological disorders with a dominant vascular component, such as stroke and Alzheimer’s disease (AD), impose a profound healthcare burden. These conditions feature heterogeneous pathobiology, driven by diverse risk factors that frequently stem from blood-brain barrier (BBB) impairment^1,2^. With over 15 million cases annually, stroke is a leading cause of death and long-term disability, and most cases are ischemic^3,4^. Early BBB disruption is a critical factor driving poor prognosis^5,7^. As a major risk factor, aging further compounds this issue through pre-existing BBB dysfunction^8–17^. Changes in blood plasma protein composition resulting from transcriptional changes in brain endothelial cells (BECs) and systemic chronic inflammation are thought to contribute to age-related BBB dysfunction^3–8^. Additionally, breakdown in BBB integrity is now recognized as a precursor to cognitive deficits, serving as an early, measurable marker for AD and dementia risk^9–12^.

The BBB is a highly selective semipermeable barrier that regulates the passage of substances between the blood and the brain. Its complex, dynamic structure comprises tight junctions (TJs) composed of proteins such as Claudin-5, Occludin, and ZO-1 in BECs, along with a low rate of caveolar transport. Supporting cells, including pericytes, astrocytes, and the capillary vascular basement membrane, contribute to the structural and functional architecture of the BBB^13^. Despite growing insights into the BBB in CNS pathology, significant knowledge gaps persist regarding the spatiotemporal mechanisms underlying its dysfunction in ischemic stroke and the development of effective targeted therapies.

The Wnt/β-catenin pathway and downstream transcription factors are critical for CNS angiogenesis, proliferation, survival, and BBB formation through stabilization and nuclear translocation of β-catenin^13^. Dysregulation of this pathway has also been implicated in stroke, AD, and MS^14–18^. More recently, small non-coding RNAs that modulate post-transcriptional gene expression, microRNAs (miRs), have emerged as important regulators of BEC function and BBB integrity^19–21^. miRs exemplify the growing recognition that non-coding RNAs possess a regulatory significance comparable to that of proteins. miRs are incorporated into Argonaute proteins to function as guide RNAs, directing the complex to target mRNAs through partial base pairing. Target recognition is partly governed by a short six-nucleotide "seed" sequence (positions 2–7), which binds complementary sites in target mRNA and results in post-transcriptional silencing by either reducing mRNA stability or suppressing protein translation^22,23^.

We and others have shown that miRs can be induced by cytokines such as interferon-β (IFN-β) and tumor necrosis factor-α (TNF-α), and that they play critical roles in modulating inflammation^24–27^. With approximately 70% of miRs expressed in the mammalian brain, these molecules are critical for maintaining CNS homeostasis and are implicated in various neurological disorders. Consequently, miRs are emerging as promising next-generation therapeutic targets for CNS diseases.

Here, we employ a non-biased anti-miR lentiviral screening strategy and identify anti-miR-23b/miR-23b as a key regulator of BBB function in BECs. Leveraging gain- and loss-of-function approaches and multi-omics, this study examines the mechanisms by which anti-miR-23b/miR-23b regulates BBB function in healthy versus ischemic stroke conditions, using 2D and 3D microfluidic *in vitro* platforms validated in *in vivo* mouse models for stroke (t-MCAO) models. Our data demonstrate that anti-miR-23b enhances BBB properties by regulating junctional proteins, transcytosis, and key signaling pathways in the brain endothelium, most notably Wnt/β-catenin, TGF-β, VEGF, and Notch. By improving BBB integrity, this approach significantly ameliorates outcomes in ischemic stroke models. These findings have high physiological and clinical relevance, positioning miR inhibitors as promising therapeutic agents.

## RESULTS

### Identification of anti-miR-23b as an enhancer of endothelial BBB properties

miRs are key regulators of brain homeostasis and are frequently dysregulated in neurological disorders, making them attractive therapeutic targets^28–32^. To identify critical negative regulators of cell-cell contact and barrier integrity, we performed an unbiased lentiviral screen using an anti-miR library targeting an array of conserved miRs, by measuring trans-endothelial electrical resistance (TEER) as a functional readout. This approach uses anti-miRs to neutralize endogenously expressed miRs, minimizing artifacts associated with ectopic overexpression. We previously validated this strategy by identifying miR-128 as a key regulator of reverse transcriptase-associated pathways, including LINE-1, telomerase, and HIV-1^33–37^.

Primary screening and validation were conducted using the following methodology. Primary mouse lung endothelial cells (ECs) were transfected with the anti-miR library to enable high-throughput loss-of-function evaluation of miR candidates. Viral expression was selected using G418 (Neomycin) resistance and validated by copGFP expression (characterized by superbright green fluorescence). Infected ECs were isolated by single-cell dilution and expanded in 96-well plates. Once EC clones reached confluence, barrier function was assessed using an electric cell-substrate impedance sensing (ECIS) instrument to measure TEER. EC wells showing a significant increase in TEER were harvested. The specific integrated anti-miR sequences were identified by cloning and validation (**Figure 1A** and **Figure S1A**). Three clones showed significantly increased TEER compared to controls (**Figure 1B**), and three anti-miRs were independently isolated from at least two different clones. Among these, anti-miR-151a and anti-miR-375 were identified as enhancers of cell junctions. We and others have previously shown that miR-151a and miR-375 play roles in tumorigenesis, and that miR-151a functions as an oncogenic miR to promote cell migration, invasion, partial epithelial-to-mesenchymal transition in non-small cell lung cancer, and tumor-associated angiogenesi^38,39^.

**Figure 1.**
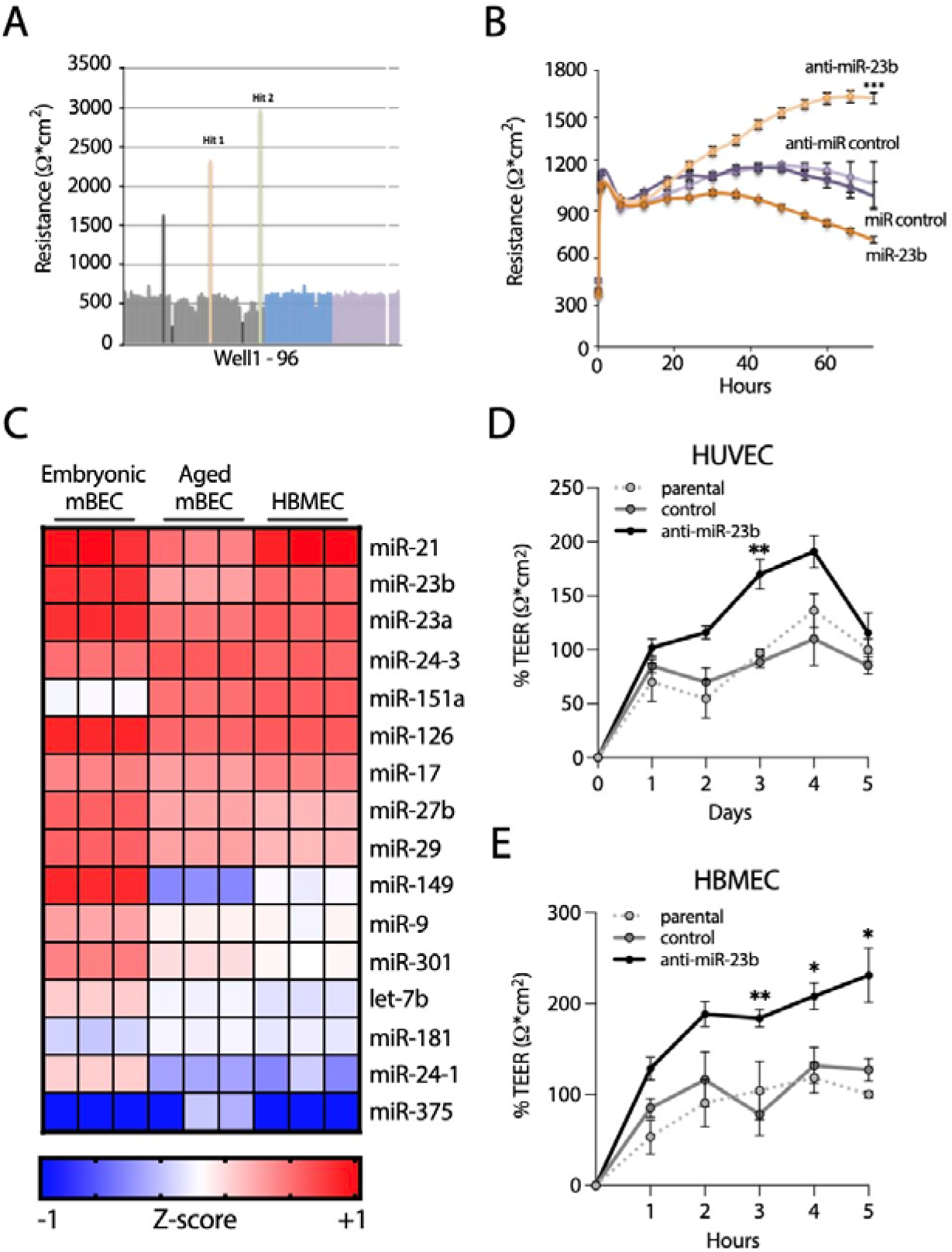
Identification of anti-miR-23b as a promoter of barrier properties in brain endothelial cells. **A**. Mouse lung endothelial cells were transduced using a pooled lentiviral-based, miR-neutralizing shRNA library, selected for puromycin resistance, and clonally expanded. Each well represents the neutralization of a single endogenously expressed miR. Cells were then analyzed by transendothelial electrical resistance (TEER) by employing an Electric Cell-substrate Impedance Sensing (ECIS) plate reader to examine paracellular barrier properties. **B.** Single high titer miR-control, miR-23b, and anti-miR-23b lentiviruses were generated and stable miR-modulated human lung endothelial cells established. TEER was measured to assess the specific and opposing effect of miR-23b and anti-miR-23b, n=12. **C.** miR expression profiling was performed by miR specific qPCR analysis of mouse brain endothelial cells (BECs) from embryonic and adult brains as well as primary human brain microvascular endothelial cells (HBMEC) and shown as a heatmap, n=6. **D.** Human HUVECs were transfected with miR control or anti-miR-23b mimics oligonucleotides, and **E.** stable HBMEC (parental, miR control, and anti-miR-23b) lines were generated. HUVEC or HBMEC cells were subjected to TEER by ECIS or EVOM3 analysis, n=4. Results shown as percent change Mean ± SEM of n=4-12 independent biological replicates, *p < 0.05, **p < 0.01, and ***p < 0.001 by ordinary-one way ANOVA at each time point.

Importantly, this screen also identified anti-miR-23b as a potent enhancer of endothelial barrier properties. Subsequently, high-titer lentiviruses encoding miR-23b and anti-miR-23b were generated. Human lung ECs were infected with lentiviruses expressing control miR, miR-23b, or anti-miR-23b and subjected to secondary TEER analysis. This confirmed that miR-23b and anti-miR-23b exert opposing effects on EC TEER measurements, establishing their role in regulating cell-cell contact and barrier integrity in lung ECs (**Figure 1C**).

To assess the relevance of the miRs identified in our screen to brain EC barrier properties, we measured the expression of several candidates in mouse and human brain endothelial cells (BECs) using miR-specific qRT-PCR. miR-375 was barely detectable in young and old mouse BECs and in primary human brain microvascular endothelial cells (HBMECs), whereas miR-151a showed low expression in young mice but robust expression in adult mice and HBMECs. In contrast, miR-23b was highly enriched across all cell types, including young and aged mouse BECs as well as HBMECs (**Figure 1C**). This aligns with previous studies that have highlighted miR-23b enrichment in the vasculature^40^. miR-23b is part of the miR-23-27-24 cluster, with two paralogous clusters (miR-23a and miR-23b) arising from ancient gene duplication events on human chromosomes 9q22 and 19p13. miR-23a and miR-23b share an identical seed sequence but differ by one nucleotide^41^, and both are expressed in the brain endothelium alongside miR-27 and miR-24 (**Figure 1C**)^40^. Given these expression patterns, miR-23b/anti-miR-23b were selected as the primary focus of this study.

To determine the functional role of anti-miR-23b in endothelial barrier integrity, we evaluated its effects on human umbilical vein endothelial cells (HUVECs) and HBMECs. In both cell models, anti-miR-23b administration significantly increased TEER compared with control cells (**Figure 1D** and **1E**). These findings indicate that miR-23b plays a critical role in modulating paracellular barrier permeability across endothelial lineages.

### Silencing miR-23b enhances the barrier properties of HBMECs by upregulating cell junction proteins

Selective BBB permeability depends on specialized adherens (AJ) and tight junctions (TJ) and their associated proteins, including VE-cadherin, Claudin-5, Occludin, and ZO-1^42,43^ (**Figure 2A**). We investigated whether miR-23b and anti-miR-23b regulate the expression of these junctional proteins in BECs.

**Figure 2.**
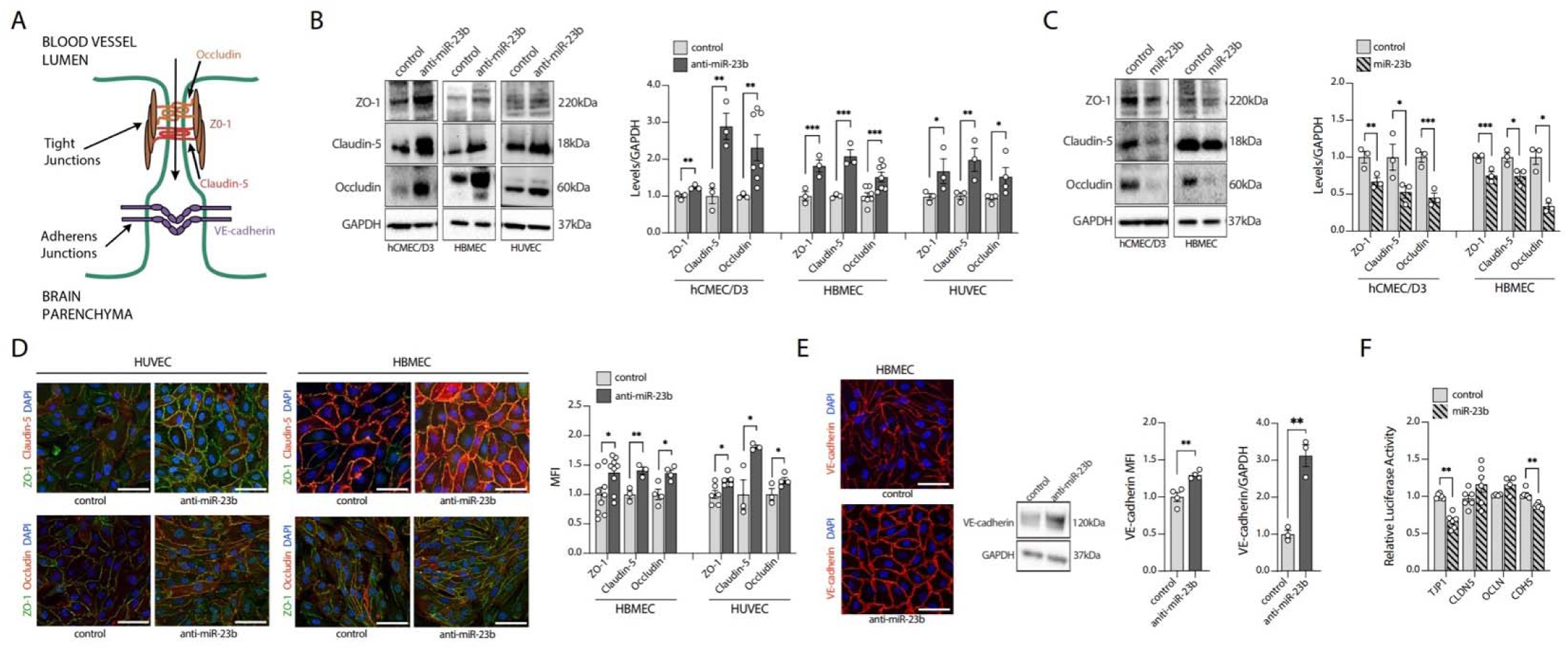
anti-miR-23b promotes barrier properties through increased expression of cell junctional proteins. **A.** Cartoon of key junctional proteins in brain endothelial cells. miR-modulated HBMEC or hCMEC/D3 cells were generated (over-expressing anti-miR-23b, miR-23b, or control miR), or HUVECs were transiently transfected with anti-miR-23b, miR-23b, or control miR oligonucleotides. Stable and transiently transduced anti-miR-23b, miR-23b, and control miR cell lines were analyzed for changes in protein levels of key tight junction proteins (TJPs; ZO-1, Claudin-5, and Occludin) by Western blot (WB) analysis. **B.** Representative WB examples of anti-miR-23b relative to control miR hCMEC/D3, HBMEC and HUVEC are shown (left panel) and quantification of WB results normalized to GAPDH are shown as relative levels, n=3 (right panel). **C.** Representative WB examples of miR-23b and controls hCMEC/D3 and HBMEC are shown (left panel) and quantifications are shown as relative levels, n=3 (right panel) (right). **D.** Representative immunofluorescence (IF) images of transduced primary HBMECs and transiently transfected HUVECs stained for ZO-1 (green) and Claudin-5 (red) or Occludin (red, left panel), and quantification of relative fluorescence levels, n=3 (right panel), scale = 50µm. **E.** Stable anti-miR-23b and control miR cell lines were analyzed for expression levels of the adherens junction protein VE-Cadherin by IF (left panel), WB (middle panel), and quantifications, n=3 (right, and far right panels), Scale = 25µm. **F.** Results of 3’UTR luciferase reporter assays for junctional is shown. HEK293 cells were co-transfected with 3’UTR containing plasmids for TJP1, CLDN5, OCLN, CDH5 or control, with miR control or miR-23b mimic oligonucleotides and luciferase activity was measured after 24hrs, n=4. Graphs are shown as Mean ± SEM (n=3-4, independent biological replicates), *p <0.05, **p < 0.01, ***p < 0.001, by two-way ANOVA for multiple comparisons and unpaired student’s t-test when comparing two groups.

We established miR-23b-overexpressing, anti-miR-23b, or control miR cell lines in both HBMECs and the human cerebral microvascular endothelial cell line (hCMEC/D3) by lentiviral transduction (routinely achieving >90% transduction efficiency). Concurrently, HUVECs were transiently transfected with miR-23b, anti-miR-23b, or control miR mimic oligonucleotides. Protein levels of ZO-1, Claudin-5, and Occludin were then assessed by Western blot (WB) analysis. Anti-miR-23b treatment markedly increased the levels of all three TJ proteins across all three endothelial cell models (**Figure 2B**), whereas miR-23b overexpression diminished their expression compared to control cells (**Figure 2C**). Building on these findings, we conducted immunofluorescence (IF) staining to determine the subcellular localization of these proteins in HUVECs and HBMECs. In both cell types, anti-miR-23b overexpression promoted robust localization of Claudin-5, Occludin, and ZO-1 at cell-cell junctions (**Figure 2D** and **Figure S2A**), indicating improved barrier integrity. Furthermore, IF and WB analyses revealed that anti-miR-23b significantly upregulated junctional VE-Cadherin levels in HBMECs compared to controls (**Figure 2D**).

To determine whether miR-23b directly targets the mRNAs of endothelial junctional proteins or regulates them indirectly, we conducted bioinformatic seed sequence analyses using TargetScanHuman 8.0 and 3’UTR luciferase reporter assays for ZO-1, Occludin, Claudin-5, and VE-Cadherin. Bioinformatic analysis revealed a conserved 8-mer seed binding site for miR-23b within the 3’UTR of TJP1/ZO-1 (Figure S2B). In contrast, no miR-23b seed binding sites were identified in the 3’UTRs of CDH5, CLDN5, and OCLN. We next conducted luciferase reporter assays to determine whether miR-23b directly binds these 3’UTR regions to suppress luciferase activity. 3’UTR-luciferase constructs of the four junctional genes were co-transfected with miR-23b or control miR mimics oligonucleotides into HEK293T cells. As shown in Figure 2F, miR-23b significantly, albeit modestly, bound to TJP1/ZO-1 and CDH5/VE-Cadherin mRNAs, whereas no binding was detected with the 3’UTRs of CLDN5/Claudin-5 or OCLN/Occludin.

These findings suggest that anti-miR-23b strengthens the endothelial paracellular barrier by upregulating tight and adherens junction proteins through both direct and indirect mechanisms. Although earlier studies established that various miRs regulate junctional proteins in cell types such as lung epithelium and endothelial cells^52,57–59^, our data demonstrate that inhibiting miR-23b in BECs is critical for promoting barrier maturation under physiological conditions.

### anti-miR-23b enhances 3D tubule formation and BBB properties of HBMECs

While traditional 2D endothelial monolayers offer limited physiological relevance for studying the BBB, 3D systems incorporating flow more accurately replicate *in vivo* vascular architecture and function. These advanced models significantly affect the expression of proteins critical to BBB integrity^44^. To better assess the impact of anti-miR-23b on BBB integrity under dynamic 3D conditions, we employed a 3D HBMEC culture model using the Mimetas 40/64-chip 3-lane OrganoPlate microfluidic system (**Figure 3A**). This platform integrates tissue-level complexity with precise environmental control, enabling reproducible, quantitative assessments of BBB properties across multiple chips. Each chip uses only ∼30,000 HBMECs, enabling cost-effective feasibility and scalability. The OrganoPlate system facilitates gravity-driven fluid flow via platform tilting, creating physiologic shear stress critical for HBMEC maturation and function. Cells are seeded into the top channel (coated with Collagen IV (Col IV)/Fibronectin (FN)), and brain capillary structures form against an extracellular matrix (ECM) gel (Col I/IV, 80:20) in the middle lane (see **Figure 3A**, left, top panel). Functional, leak-tight 3D BBB tubules are visualized by performing barrier integrity (BI) assays using fluorescent dextran tracers of various molecular weights and colors, which are perfused into the upper channel to confirm the selectivity and maturity of HBMEC tubules. BI assays using a green 150KDa dextran tracer (**Figure 3A**, right panel) and 3D expression of TJ proteins (Claudin-5 and ZO-1) demonstrate functional brain capillary-sized tubules (**Figure 3B**).

**Figure 3.**
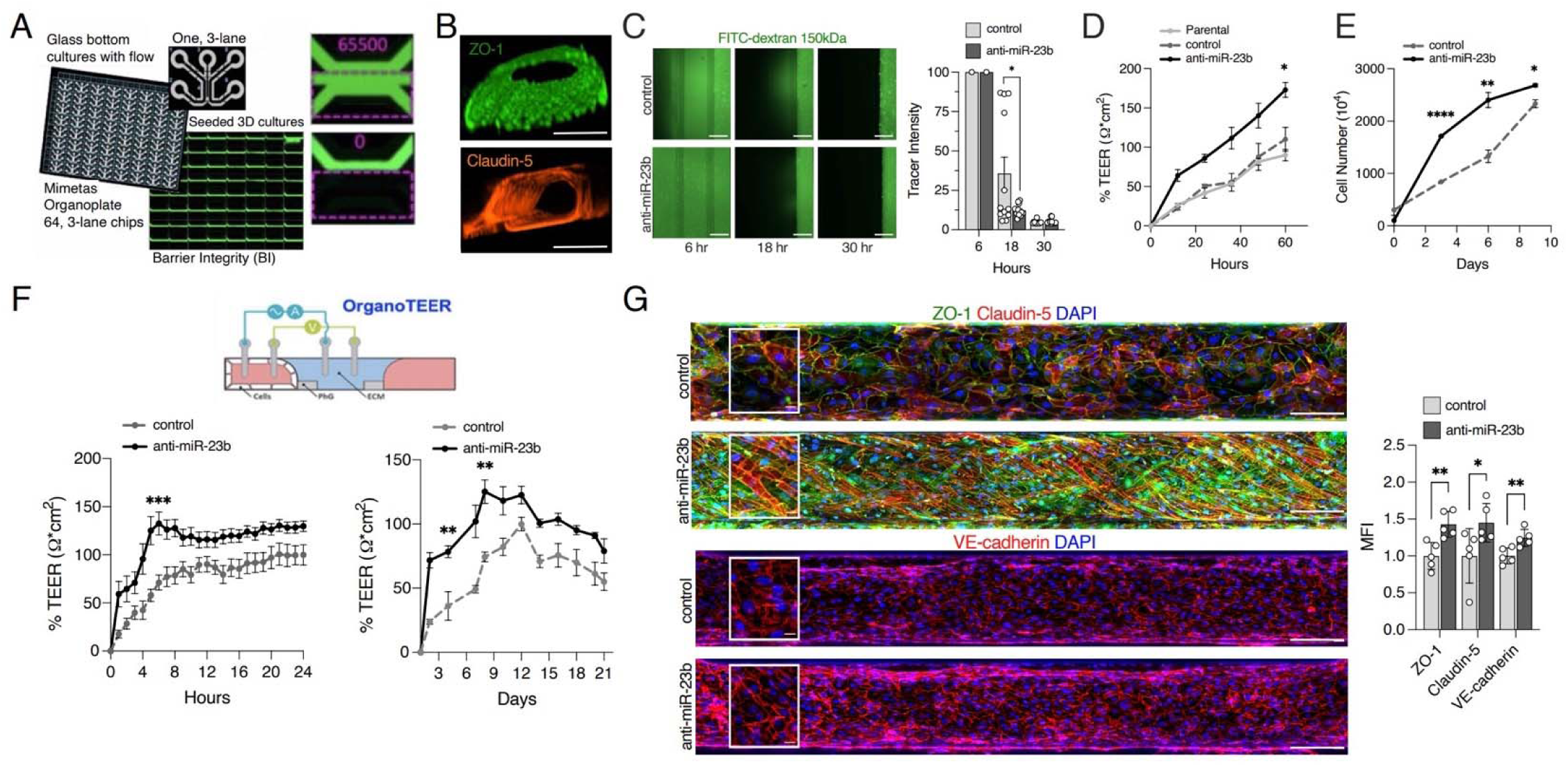
anti-miR-23b accelerates 3D brain endothelial tubule formation and strengthen their barrier properties. **A.** Outline of the Mimetas organ-on-a-chip plate. Perfused 3D tubules are established in 40 or 64 3-lane chip cultures using the Mimetas platform. The top channel is seeded with HBMEC against the ECM (middle channel), allowing for 3D tubule formation of brain capillary size, and the bottom channel can be seeded with CNS cells. A tight tubule only shows tracer in the top channel of the 3-lane culture chip, whereas a leaky tubule will show tracer in the middle channel. Barrier integrity (BI) analysis is performed using different MW-sized tracers. Shown are intact HBMEC tubules in a 64-chip plate using a 150KDa FITC-dextran (green) tracer, and quantification of BBB permeability of a ‘no cell’ chip culture showing complete leakiness (top) and an HBMEC chip showing no permeability, representing a functional BBB (bottom). **B.** Intact 3D HBMEC tubule formation is visualized by ZO-1 and Claudin-5 staining. **C.** The timing of tubule formation was examined using anti-miR-23b and miR control HBMECs by BI analysis at 6, 18, and 30 hours, n=3. Scale = 250µm. **D.** 3D TEER measures of parental, control miR and anti-miR-23b HBMECs was recorded, n=12. **E.** The proliferation potential of anti-miR-23b and control HBMECs was determined by cell counting of triplicate cultures, using a cell coulter counter, n=5. **F.** Schematic of the 3D OrganoTEER equipment is shown (top, approved cartoon modification from Mimetas website) and plots of kinetic analysis of HBMEC tubule establishment (left panel) and long-term tubule resilience (right panel) were built from the TEER measurements, n=12-16. **G.** Shown are IF staining for key junctional proteins [ZO-1 (green), Claudin-5 (red) or VE-Cadherin (red), and nuclear stain (DAPI, blue)] in anti-miR-23b and control 3D HBMEC tubules. Representative images of the full top channel tubule and magnified images (left panels) and quantifications are shown (right panel), n=3, Scale = 250µm and 25µm. All graphs are shown as Mean ± SEM, n= 3-16 independent biological replicates, *p <0.05, **p < 0.01, ***p < 0.001, ****p < 0.0001, by two-way ANOVA for multiple comparisons and multiple unpaired t-tests at each time point for TEER measurements.

To determine how anti-miR-23b affects 3D vascular formation and barrier function in HBMECs, we compared anti-miR-23b-expressing HBMECs to control miR cells in the 3D assay. Both groups formed morphologically intact tubules; however, anti-miR-23b significantly accelerated the formation of leak-tight HBMEC tubules, and tighter paracellular barrier function was observed as early as 18 hours post-seeding. In contrast, control tubules remained permeable at 18 hours (**Figure 3C**). This observation was also supported by 3D TEER measurements with the OrganoTEER, which showed significantly higher resistance in anti-miR-23b cultures than in parental and control-miR HBMECs (**Figure 3D**). Next, we tested whether anti-miR-23b affects HBMEC cell proliferation, observing a modest but significant increase in cell proliferation potential in anti-miR-23b HBMECs, compared to control HBMECs, as previously reported^63,64^, likely contributing to the anti-miR-23b HBMECs’ enhanced barrier properties (**Figure 3E**).

Additionally, we tracked TEER during the first 24 hours of tubule development to assess tubule establishment (**Figure 3F**, top). anti-miR-23b significantly accelerated the formation of functional tubules compared with controls (**Figure 3E**, left panel). Moreover, anti-miR-23b strengthened tubule resistance over three weeks in culture (**Figure 3F**, right panel).

Finally, we examined expression levels of key junctional proteins by 3D IF staining for ZO-1, Claudin-5, and VE-Cadherin using a C10 confocal imaging plate reader, demonstrating intact, lumenized 3D tubules in anti-miR-23b cultures with significantly elevated expression of ZO-1, Claudin-5, and VE-Cadherin compared to controls (**Figure 3G**).

Confirming our 2D data (**Figure 2**), these findings demonstrate that anti-miR-23b enhances HBMEC barrier function and maturation in a physiological 3D context. Inhibiting miR-23b accelerated and strengthened the formation of a 3D microfluidic BBB model, with robust junctional protein expression and sustained structural integrity.

### Inhibiting miR-23b enhances transcellular barrier properties by downregulating PLVAP

In addition to TJ integrity, low rates of transcellular transport are essential for maintaining BBB function^42,45^. Dysregulation of transcellular transport has been recognized as an early contributor to barrier breakdown following pathological insults such as stroke^46,47^. To assess whether anti-miR-23b modulates transcytosis in BECs, we examined levels of several proteins that regulate transcellular transport in anti-miR-23b-expressing and control HBMECs and hCMEC/D3 cell lines (see **Figure 4A**). WB analysis revealed that anti-miR-23b significantly downregulated Plasmalemma Vesicle-Associated Protein (PLVAP), a marker of fenestrated ECs (**Figure 4B**). In contrast, Caveolin-1 (Cav-1) and Caveolin-2 (Cav-2, not shown) protein levels were not altered by anti-miR-23b expression (**Figure 4B**). We performed a transcellular uptake assay using fluorescently labeled albumin (albumin-Alexa594, red) to determine the functional consequences of PLVAP downregulation. After one hour of exposure, anti-miR-23b HBMECs showed a marked reduction in intracellular albumin-Alexa594 signal compared to control cells, as shown in a panel of 3D HBMEC chip cultures (**Figure 4C**), and in a representative 3D full tubule image at higher magnification, indicating reduced uptake (**Figure 4D**, left panel). Quantification confirmed a significant decrease in albumin uptake in anti-miR-23b cultures (**Figure 4D**, right panel). To independently validate this observation, we performed IF imaging of PLVAP and DAPI in anti-miR-23b versus control HBMEC cultures. anti-miR-23b expression led to a modest but statistically significant reduction in PLVAP protein levels (**Figure 4E**), consistent with the WB findings. Therefore, in BECs, anti-miR-23b acts as a dual stabilizer of the BBB by reducing transcellular transport, via PLVAP downregulation, and enhancing the structural integrity of junctional proteins.

**Figure 4.**
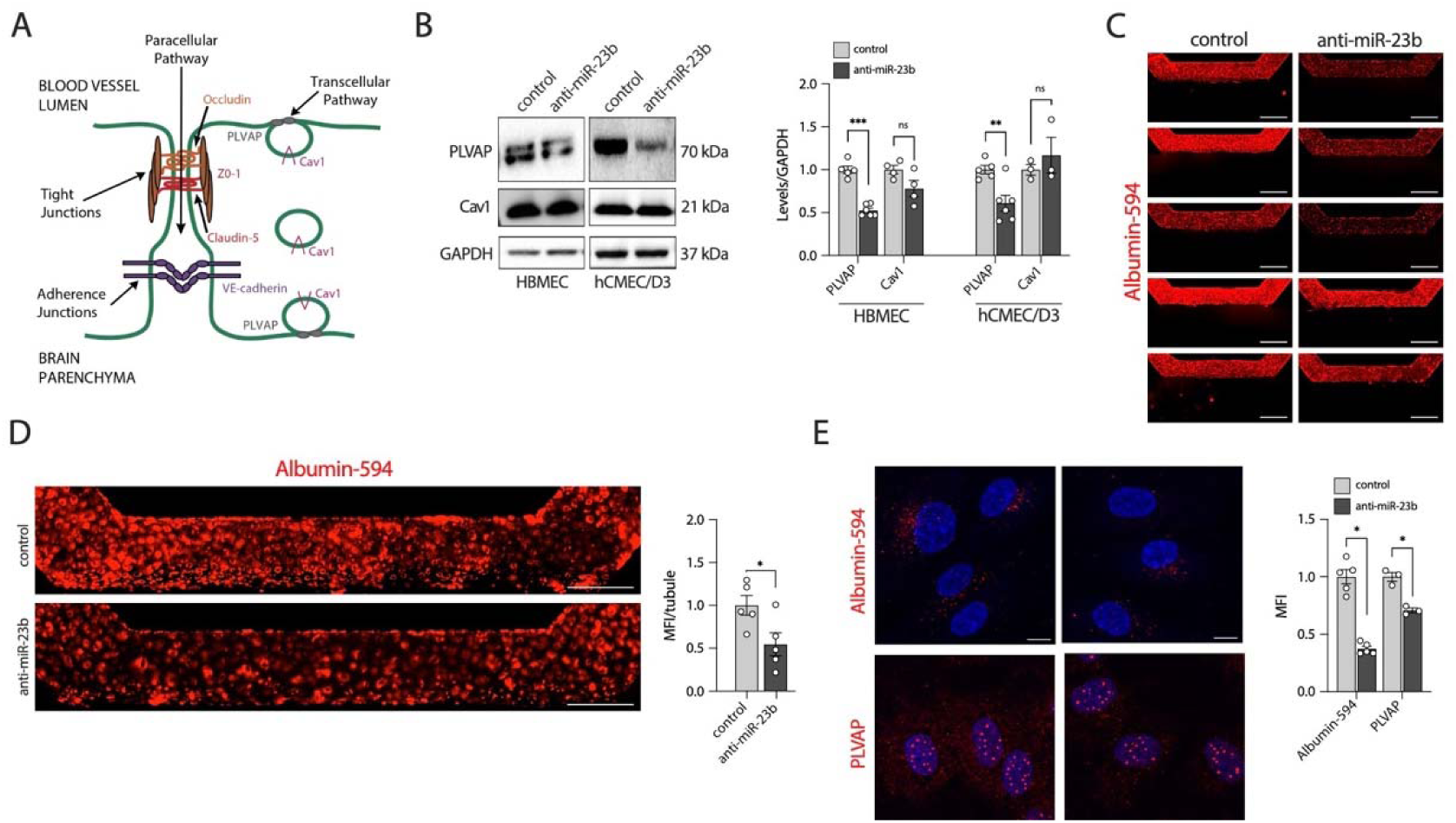
anti-miR-23b strengthen barrier properties by reducing transcytosis and downregulating PLVAP protein levels. **A.** Cartoon of brain endothelial cell junctional proteins and transcellular transport, which is regulated by PVLAP. **B.** miR-control or anti-miR-23b overexpressing HBMEC or hCMEC/D3 cells were analyzed for changes in expression levels of Caveolin-1 (Cav1) and PLVAP by Western blotting. Representative examples are shown (left panel), and quantification of results normalized to GAPDH as relative levels, n=3 (right panel). **C, D.** Transcellular uptake of labelled Albumin-594 (red, readout of transcellular transport), were assessed in control or anti-miR-23b HBMECs. **C**. Albumin-594 transcytosis is shown in multiple chip cultures, n=5. Scale = 500µm. **D.** A representative images of a single 3D tubule (left panel) and quantification of albumin transcytosis (right panel), n=5. Scale = 25µm. **E.** Confocal images of PLVAP/DAPI staining of anti-miR-23b or controls (left panel) and quantifications, n=3 (right panel), Scale = 12.5µm. Graphs are shown as Mean ± SEM, n=3-5 independent biological replicates, *p <0.05, **p < 0.01, ***p < 0.001, ****p < 0.0001, by two-way ANOVA for multiple comparisons and by unpaired two-tailed Student’s t-test.

### Inhibiting miR-23b enhances barrier properties through Wnt/β-catenin-dependent and - independent mechanisms

Wnt/β-catenin signaling plays a central role in promoting and maintaining BBB integrity^42,48,49^. CNS injury triggers reactivation of Wnt/β-catenin signaling, which contributes to vascular repair mechanisms^55^. Given this pathway’s essential role in preserving the BBB, pharmacological or genetic modulation of Wnt/β-catenin represents a promising therapeutic approach in preclinical models of neurological disorders, such as ischemic stroke^14–18^. To determine whether miR-23b regulates the Wnt/β-catenin pathway, we first analysed protein levels of β-catenin and its transcriptional coactivator LEF-1 in anti-miR-23b-expressing versus control ECs. WB analysis revealed that anti-miR-23b significantly increased the protein levels of β-catenin and LEF-1 in hCMEC/D3, HUVEC, and HBMEC lines (**Figure 5A**). We next corroborated these findings using IF in 2D cultures and 3D HBMEC tubules. There was increased expression of β-catenin and LEF-1 in anti-miR-23b HBMECs compared to controls in both 2D and 3D models (**Figure 5B** and **Figure S4A-B**).

**Figure 5.**
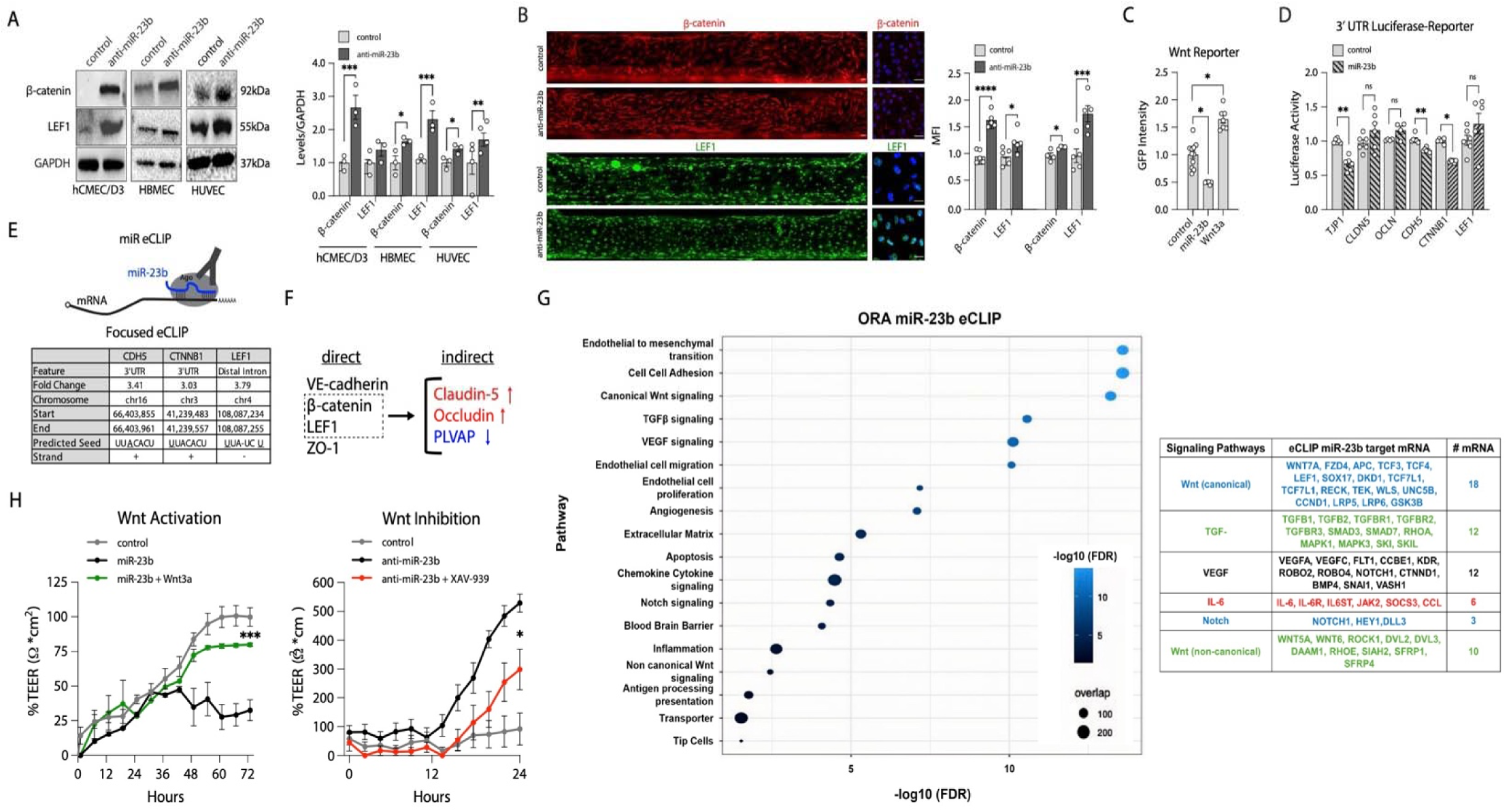
anti-miR-23b enhances barrier properties through Wnt/β-catenin dependent and independent mechanisms. **A.** Levels for β-catenin and LEF-1 proteins were examined by Western blot analysis in anti-miR-23b or control HBMECs, hCMEC/D3 transduced cells, and in control or anti-miR-23b mimic oligonucleotide transfected HUVECs. Results are shown as representative images (left panels) and quantified, n=3 (right panels). **B.** Representative IF images of stable transduced HBMEC (control and anti-miR-23b) stained for β-catenin (red) or LEF-1 (green) in 3D tubules (left panel), magnification of β-catenin and LEF-1 expression in HBMECs in 2D (middle panel) and quantifications (right panels), n=3. Tubule scale = 25µm. Magnified image of β-catenin and LEF-1 scale = 50µm and 30µm. **C.** Wnt-GFP reporter assays using HEK293 cells transfected with M38-TOP-dGFP reporter plasmids with miR control or miR-23b mimic oligonucleotides, using Wnt3a as a positive control, n=6. **D.** 3’UTR-luciferase assays were performed for β-catenin (CTNNB1) and LEF-1 by transfecting HEK293 cells with 3’UTR-luciferase constructs for either CTNNB1, LEF-1 or control with either control or miR-23b mimic oligonucleotides, n=7. Shown are results of junctional proteins and β-catenin and LEF-1. **E.** miR enhanced crosslinking and immune-purification (miR-eCLIP) sequencing analysis, was performed for miR-23b, see cartoon (top panel). Control, miR-23b and anti-miR-23b overexpressing HBMEC lines were generated, their barrier phenotype was validated by TEER analysis and miR eCLIP-seq was performed. The Table shows details of miR-23b direct binding to VE-Cadherin (CDH5), β-catenin (CTNNB1) and LEF-1 mRNA. Underlined nucleotides in predicted seed, highlight mismatches. **F.** Cartoon representation of likely miR-23b effects in BECs (direct or indirect) on VE-Cadherin, β-catenin, LEF-1, ZO-1 and Claudin-5, Occludin and PLVAP. **G.** Global, unbiased over-representation analysis (ORA) was performed on HBMEC eCLIP results, showing enriched pathways as listed (left panel), and examples of miR-23b mRNA targets for the top signaling pathways (Table, right panel). **H.** To assess the contribution of Wnt/β-catenin signaling of the miR/anti-miR-23b effect in BECs, rescue experiments were performed. To examine the effect of Wnt activation, control, miR-23b or miR-23b HBMEC with Wnt3a (Wnt agonist) were challenged to TEER analysis, rescue effect shown in green (left panel). To examine the effect of Wnt inhibition, control, anti-miR-23b or anti-miR-23b HBMEC with XAV-939 (Wnt inhibitor) were challenged to TEER analysis, rescue effect showed in red (right panel), n=5. Graphs are shown as Mean ± SEM, n=3-7 independent biological replicates, *p <0.05, **p < 0.01, and ****p < 0.0001, by two-way ANOVA for multiple comparisons, one-way ANOVA when analyzing one variable, unpaired two-tailed Student’s t-test for two groups and multiple unpaired t-tests when analyzing TEER measurements.

Next, we performed Wnt-GFP reporter assays to assess whether miR-23b affects Wnt/β-catenin activity. Briefly, HEK293 cells were transfected with the M38-TOP reporter along with control or miR-23b mimic oligonucleotides, and Wnt3a served as a positive control for pathway activation. The assay revealed a significant reduction in Wnt-GFP reporter activity in the presence of miR-23b, demonstrating that miR-23b inhibits Wnt/β-catenin signaling (**Figure 5C**).

To determine whether miR-23b directly targets β-catenin and LEF-1 mRNA, 3’UTR luciferase assays were performed as previously described. The results showed a modest but significant decrease in β-catenin 3’UTR reporter activity, with no direct binding to LEF-1 mRNA. Reporter assay results for the junctional proteins and for β-catenin/LEF-1 interactions are shown in **Figure 2F** and **Figure 5D**, respectively.

Because miRs often bind with imperfect seed complementarity and can target mRNAs outside the 3’ UTR, we used miR eCLIP-sequencing to examine in-cell miR-mRNA interactions. By capturing chimeric RNA reads formed within the AGO2 complex, this approach directly maps miR-23b-mRNA binding sites—surpassing the limitations of computational predictions and reporter assays (**Figure E**, top panel). To define the direct targets of miR-23b, control, miR-23b-overexpressing, and anti-miR-23b-overexpressing HBMECs were subjected to miR-eCLIP-Seq. Focused analysis of junction-related and Wnt/β-catenin pathway genes, including CDH5/VE-Cadherin, CLDN5/Claudin-5, OCLN/Occludin, TJP1/ZO-1, CTNNB1/β-catenin, and LEF1, revealed that miR-23b directly binds to CDH5, CTNNB1, and LEF1 mRNAs, showing more than 3-fold enrichment compared to controls (**Figure 5E**, Table, **Figure S4C** and **Extended Data Set 1**). Interestingly, these binding regions included non-perfect seed sequences, specifically within the 3’ UTRs of CDH5 and CTNNB1 and the distal intron of LEF1 (**Figure 5E**). These miR-23b targets were successfully validated in hCMEC/D3 cells, lymphatic endothelial cells (LECs), or both (data not shown).

These combined results establish that miR-23b directly targets the mRNAs of CDH5, CTNNB1, and LEF-1. miR-23b may also bind TJP-1 mRNA in certain cellular contexts, based on 3’UTR-luciferase and bioinformatic analyses. However, Claudin-5 and Occludin appear to be indirect targets of miR-23b, likely regulated via the Wnt/β-catenin signaling pathway. As illustrated in the cartoon, anti-miR-23b modulates this pathway to increase Claudin-5 and Occludin expression while reducing PLVAP (**Figure 5F)**.

Next, we performed an unbiased analysis of the miR-23b eCLIP-seq datasets to examine global effects of miR-23b/anti-miR-23b in HBMEC, identifying prominently altered pathways and further assessing the role of Wnt/β-catenin signaling. These analyses established that, in addition to CTNNB1 and LEF-1, miR-23b targets several other key mRNAs in the canonical Wnt/β-catenin pathway, including WNT7A, FZD4, LRP5, LRP6, APC, TCF3, TCF4, and DKK1.

As shown in the Over-Representation Analysis (ORA) dot plot, miR-23b targets are enriched for numerous key signaling pathways, including the TGF-β pathway (TGF-β1, TGF-β2, TGF-βR1, TGF-βR2), VEGF (VEGFA, VEGFC, FLT1), Notch (NOTCH1, HEY1, DLL3), IL-6 (IL-6, IL-6R, IL6ST), and the non-canonical Wnt/β-catenin pathway (WNT5A, ROCK1, DVL2, DVL3, DAAM1) (**Figure 5G**, left and **Figure 5G** Table, right). The complete miR eCLIP target list is available as **Extended Data Set 1.**

Collectively, these findings indicate that anti-miR-23b elicits robust, multifaceted effects on BBB integrity. Specifically, the canonical Wnt/β-catenin pathway promotes structural maturation by enhancing tight junction formation and suppressing vesicular transport. Concurrently, non-canonical Wnt signaling modulates cell polarity and stress responses, and both pathways act in concert to fortify barrier stability. anti-miR-23b enhances BBB properties by promoting maturation of the neurovascular unit, specifically by increasing TJ protein expression via TGF-β and the MEK/ERK pathway. Notch signaling is a critical, conserved regulator of BBB integrity and cerebrovascular homeostasis, whereas VEGF and IL-6 signaling typically increase BBB permeability, particularly following injury. However, under precise, low-dose conditions, VEGF and IL-6 can promote protective angiogenesis and neurogenesis.

The ORA analysis further revealed that miR-23b targets numerous mRNAs and proteins critical for cell-cell adhesion (CDH5, CTNNA1, PECAM1, JAM3), angiogenesis (NOTCH1, VEGFA, SNAI1, SERPINE1), BBB function (CLDN12, JAM3, CDH5, FOXP1, FOXO1), and the extracellular matrix (ECM) (COL4A1, COL4A2, LAMB1, LAMA2, LAMC1, ITGA2, ITGB3). Together, these effectors are essential for maintaining BEC function and BBB integrity in health and disease.

Finally, to examine Wnt/β-catenin-dependent and -independent mechanisms of miR/anti-23b-related effects on BBB properties, we performed dual rescue experiments. First, we assessed Wnt/β-catenin activation by culturing miR control or miR-23b HBMECs in the presence or absence of Wnt3a, a Wnt agonist, and measured the effect on 3D TEER. If the Wnt/β-catenin pathway is required for the anti-miR-23b protective phenotype, the Wnt agonist would be expected to inhibit the miR-23b-induced loss of barrier integrity (**Figure 5H**, left panel in green). Second, we evaluated Wnt/β-catenin inhibition by applying a specific Wnt/β-catenin inhibitor (XAV-939) to control or anti-miR-23b HBMEC cultures and measured effects on TEER. If Wnt/β-catenin signaling is required for the protective BBB phenotype, the inhibitor should attenuate the anti-miR-23b-mediated rescue of barrier properties (**Figure H**, right panel in red).

The rescue experiments show that the miR/anti-miR-23b effect is partly dependent on Wnt/β-catenin signaling. Anti-miR-23b-enhanced TEER was significantly but partially reduced by the Wnt inhibitor (XAV-939), and miR-23b-induced loss of TEER was partially rescued by Wnt3a. Together, these orthogonal gain- and loss-of-function manipulations establish Wnt/β-catenin signaling as a partial but necessary mediator of the protective barrier function of anti-miR-23b.

Collectively, our findings demonstrate that miR-23b and anti-miR-23b regulate barrier properties by modulating the Wnt/β-catenin signaling pathway, specifically by targeting β-catenin, LEF1, and key downstream effectors in BECs. Furthermore, by orchestrating this pathway alongside the TGF-β, Notch, VEGF, and IL-6 networks, anti-miR-23b reinforces barrier integrity, whereas miR-23b diminishes it.

Future studies will determine the specific contributions of these pathways to miR-23b’s role in BEC and BBB biology. However, our findings establish that miR-23b and anti-miR-23b are important regulators of Wnt/β-catenin signaling in BECs. By enhancing β-catenin, LEF1, and downstream effector proteins, anti-miR-23b drives a protective, barrier-enhancing phenotype. Furthermore, anti-miR-23b affects additional key signaling cascades, including TGF-β, Notch, VEGF, and IL-6, which are inversely regulated by miR-23b. Future analysis will delineate the specific contribution of each pathway to the overall modulation of BEC and BBB biology by the miR-23b system.

### Unbiased analysis of miR/anti-miR-23b effects on the transcriptome and proteome of HBMEC

To provide a global unbiased view of genes, proteins and pathway affected by miR-23b/anti-miR-23b in HBMECs, we next generated transcriptomic and proteomic profiles comparing control, miR-23b, and anti-miR-23b-overexpressing HBMECs to examine global transcriptional and translational changes elicited by miR-23b and anti-miR-23b. In brief, miR/anti-miR/control miR HBMECs were generated, expanded, and the barrier phenotype was validated by TEER analysis. Cells were harvested, RNA was subjected to RNA sequencing, and protein was subjected to proteomic analysis from parallel cultures.

First, principal component analysis (PCA) of control vs miR-23b and anti-miR-23b HBMEC gene profiles was performed, demonstrating separation of the 3 conditions (**Figure 6A**). One anti-miR-23b sample was identified as an outlier. The outlier sample was included in all differential gene expression (DGE) analyses but was excluded from the heatmaps shown in **Figure 6D-G**. Heatmaps including all samples are shown in **Figure S5.2.**

**Figure 6:**
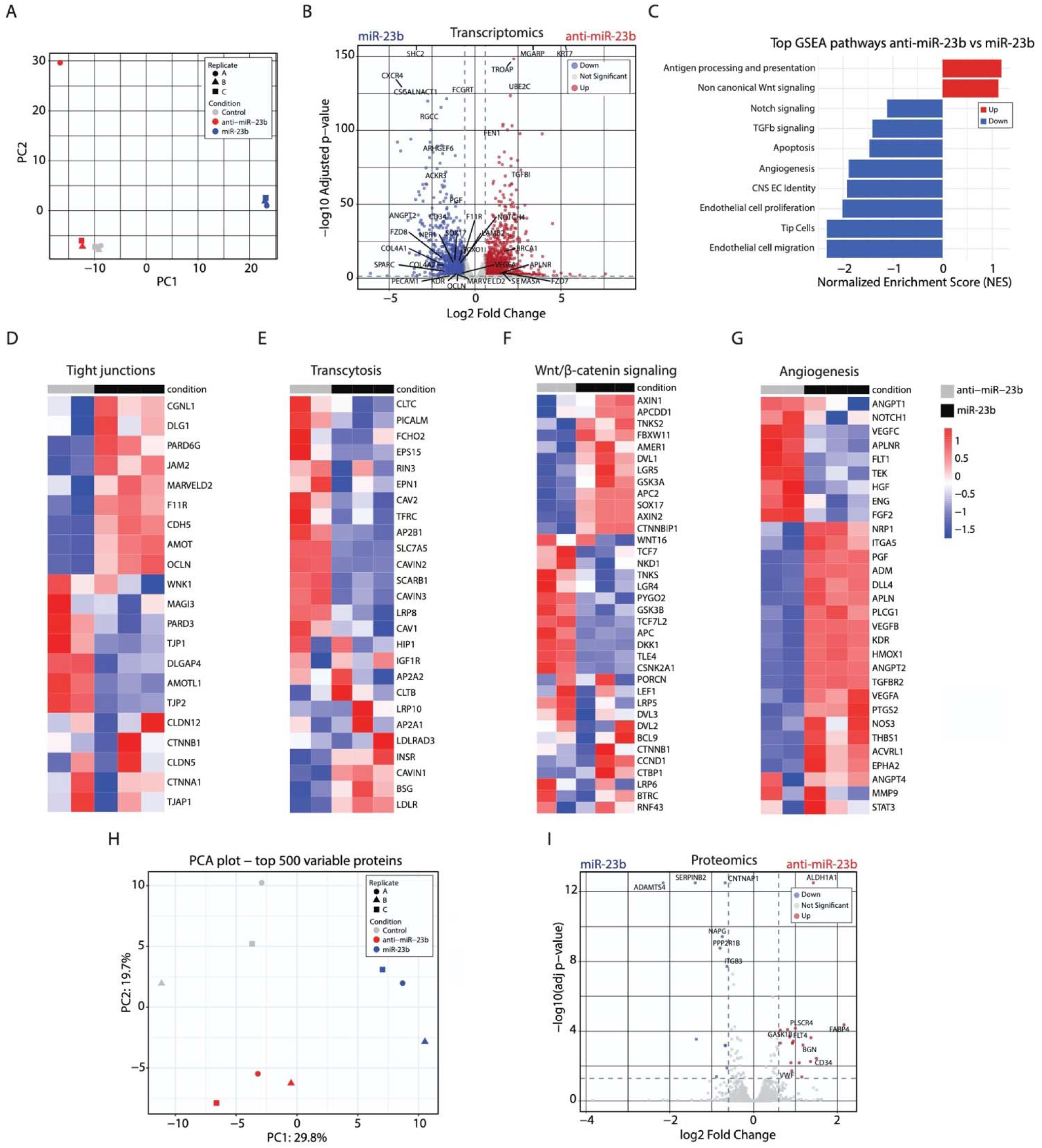
anti-miR-23b and miR-23b HBMEC transcriptome and proteome analysis demonstrate changes in genes and proteins regulating angiogenesis and barrier properties. **A.** Principal component analysis (PCA) plot illustrating control vs miR-23b versus anti-miR-23b HBMEC gene profiles. One anti-miR-23b sample was determined. The outlier sample was included in all DEG analysis but was removed from the shown heatmaps**. B.** Volcano plot highlighting upregulated and downregulated BBB-specific genes in anti-miR-23b compared to miR-23b HBMECs. All listed genes are significant (Padj<0.05; |logFC|>0.25). **C.** Gene Set Enrichment (GSEA) analysis pathways in anti-miR-23b compared to miR-23b HBMEC. Bars indicate −log(false discovery rate) values. **D** - **G,** Heatmap of BBB-specific genes (TJ [tight junction] proteins, receptor-mediated transcytosis, canonical Wnt/β-catenin signaling and Angiogenesis between anti-miR-23b and miR-23b HBMECs (one samples was an outlier, see PCA plot in A, and was removed, but Heatmaps with all samples are shown in Figure S6.2). The scale bar is log(Z score), where dark colors indicate a high Z score and light colors indicate a low Z score. **H.** PCA plot illustrating control vs miR-23b versus anti-miR-23b HBMEC protein profiles. **I.** Volcano plot highlighting upregulated and downregulated BBB-specific proteins in anti-miR-23b compared to miR-23b HBMECs. All listed proteins are significant (Padj<0.05; |logFC|>0.6).

Next, differential gene expression analysis was run, and the results visualized in a volcano plot (**Figure 6B**), showing the upregulated and downregulated genes including some of the BEC-specific genes in anti-miR-23b compared to miR-23b (**Figure 6B**). These analyses show that miR-23b upregulates genes related to angiogenesis (e.g., CXCR4, ACKR3, ANGPT2, FZD8, CD34, SPARC, KDR/VEGFR2, PGF, VEGFA), promoting endothelial cell migration, angiogenesis, vascular destabilization, and permeability, whereas anti-miR-23b upregulates some BBB-specific genes (e.g., TGFB1, NOTCH4, BRCA1, APLNR, FZD7, PGARP, KRT7), promoting vascular stability and BBB maintenance. These findings align with the concept that downregulation of angiogenic genes strengthens the BBB by shifting endothelial cells from a proliferative, high-permeability state to a quiescent, stable state, thereby allowing tight junction maturation. Reducing angiogenic signaling, particularly VEGF and components of the Wnt pathways, prevents the formation of "leaky" vessels and enhances barrier integrity.

The results from the DGE analysis were then used for Gene Set Enrichment Analysis (GSEA) to assess pathways regulated in anti-miR-23b samples compared with miR-23b samples. anti-miR-23b HBMECs showed upregulated genes relevant to non-canonical Wnt signaling and antigen processing and presentation, whereas miR-23b samples showed upregulated genes relevant to Notch and TGF-β signaling, cell migration, angiogenesis, and CNS/EC cell identity (**Figure 6C**). These results strengthen the finding that miR-23b enhances angiogenesis and CNS/EC cell identity, whereas anti-miR-23b enhances Wnt signaling and antigen processing and presentation, which is critical for maintaining barrier stability and immune surveillance at the BBB. GSEA analysis of control versus miR-23b DEGs confirmed that miR-23b promotes angiogenesis, whereas analysis of anti-miR-23b versus control confirmed suppression of angiogenic pathways and upregulation of non-canonical Wnt signaling (**Figure S5.1A-B**).

Additionally, we generated heatmaps of BBB-specific genes (TJ proteins, receptor-mediated transcytosis, canonical Wnt/β-catenin signaling, and angiogenesis) in anti-miR-23b and miR-23b HBMECs (**Figure 6D-G**), to demonstrate specific genes that are either up- or downregulated by miR-23b and anti-miR-23b. Anti-miR-23b treatment upregulated a subset of BBB-transcriptome related to tight junctions, transcytosis, transporters and extracellular matrix proteins (**Figure 6D & E, Figure S5.1C & D** and **Extended Data Set 2)**. In addition, a subset of the Wnt/β-catenin signaling genes are upregulated in anti-miR-23b samples, whereas numerous angiogenesis genes are downregulated by anti-miR-23b compared to miR-23b (**Figure S5.1C-F, Figure S5.2A-H** and **Extended Data Set 2).** Finally, anti-miR-23b upregulated a subset of genes related to CNS/EC identity, and non-canonical Wnt signaling compared to miR-23b (**Figure S5.1E & F**). All heatmaps, including the outlier sample, are included in **Figure S5.2A-H**, along with gene lists of miR/anti-miR-23b and control vs miR or anti-miR-23b in **Extended Data Set 2)**.

Because miRs affect both mRNA expression and protein translation, proteomic analysis was performed on parallel HBMEC samples. Proteomic PCA plots between control vs miR-23b vs anti-miR-23b HBMEC protein profiles demonstrated clear separation among conditions (**Figure 6H**). Volcano plots visualizing the up- and down-regulated proteins in anti-miR-23b compared to miR-23b (**Figure 6I** and **Extended Data Set 3**), confirmed upregulation of several proteins (e.g., SERPINEB2, ADAMTS4, TGFBI, ITGB3) known to regulate barrier integrity and promote angiogenesis and inflammation. In contrast, very few proteins were upregulated by anti-miR-23b overexpression (e.g., MGP, CD34, ALDH1A1, PLSCR4, FGF2) (**Figure 6I and Extended Data Set 3**). Some of these proteins support BBB integrity, stabilize vascular structures, and are neuroprotective and anti-inflammatory. Control vs miR-23b or control vs anti-miR-23b volcano plots also confirmed similar findings (**Figure S5.3A-B).** The proteomic analysis validated some molecules identified as miR/anti-miR-23b regulated genes in the transcriptomic analysis (ITGB3, CNTNAP1, NAPG, SERPINB2 and ADAMTS4). This confirms that changes in these specific transcripts are reflected in corresponding changes at the protein level.

In summary, transcriptomic and proteomic analyses reveal opposing roles for miR-23b and anti-miR-23b in BECs: miR-23b potently induces angiogenesis-specific genes and proteins, whereas anti-miR-23b reverses this effect, upregulating a subset of BBB-specific markers. Moreover, several pro-angiogenic pathways (e.g. Notch, TGF-β) were suppressed in anti-miR-23b samples to promote a functional BBB. Integrating eCLIP-seq results with transcriptomic and proteomic signatures demonstrates that miR-23b/anti-miR-23b regulates a subset of the large network of BBB genes and functions, with critical implications for endothelial health and disease.

### anti-miR-23b reduces BBB deficits in a 3D in vitro stroke-like model

Ischemic stroke induces BBB dysfunction through a cascade of cellular events, including serum protein leakage, immune cell infiltration, and inflammation, that exacerbate neuronal damage^50–52^. Even subtle increases in BBB permeability can lead to hippocampal injury and cognitive declin^53–57^. However, the underlying mechanisms of chronic BBB dysfunction after ischemic injury remain incompletely understood. We used our established 3D HBMEC culture platform to model ischemic stroke *in vitro* using the oxygen-glucose deprivation (OGD) paradigm. Baseline cultures of both anti-miR-23b and control HBMECs were established as leak-tight 3D tubules before OGD exposure (days 7–10). 3D HBMEC tubules were then exposed to hypoxia (1% O₂), glucose deprivation, and static flow for 6 hours, then returned to normoxic, glucose-replete conditions with restored flow (**Figure 7A**). This setup enabled us to assess both "acute injury”, and "longer-term recovery" mechanisms related to BBB integrity. TEER analysis showed that while both groups exhibited a decline in barrier function during OGD, the loss was significantly less severe in anti-miR-23b cultures. Furthermore, anti-miR-23b cultures showed enhanced recovery following stroke-like injury (**Figure 7A**), suggesting a protective role for anti-miR-23b in injury resistance and barrier repair. BI assays further confirmed reduced BBB leakiness in anti-miR-23b cultures at both early (5 and 8 hours) and later timepoints (3 and 14 days) post-stroke-like conditions (**Figure 7B**).

**Figure 7.**
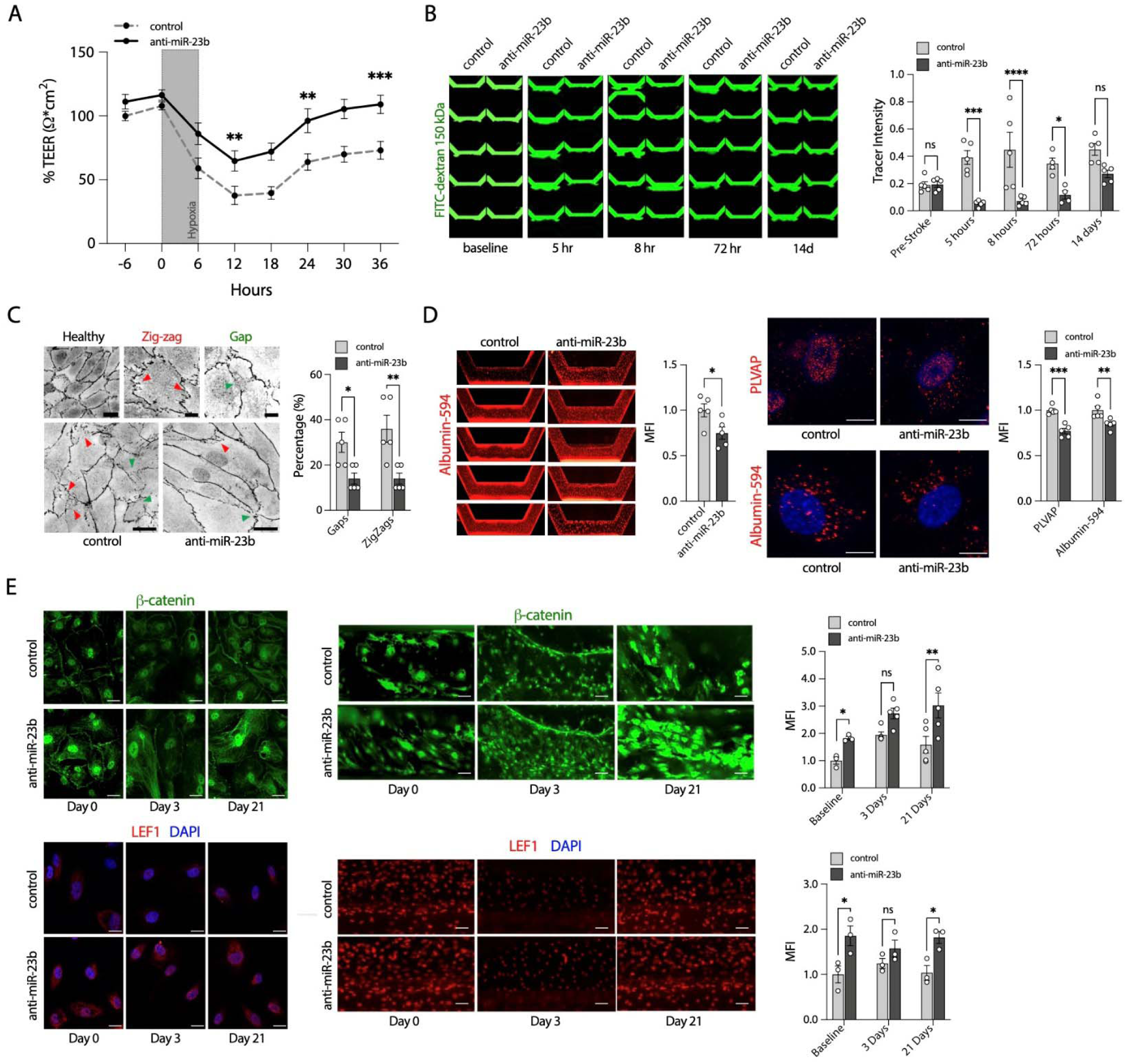
anti-miR-23b reduces BBB deficits in a 3D *in vitro* stroke model. **A.** anti-miR-23b or control HBMECs were seeded in the 3D Mimetas device. When 3D tubules were intact, cultures were exposed to *in vitro* stroke-like conditions (1% hypoxia, no glucose media, no/low flow). Continuous 3D TEER analysis shows partial loss of barrier function over time (n=10 chip cultures/condition). After 6 hours of *in-vitro* stroke-like conditions, cultures were rescued (moved to normoxia) with glucose, and perfusion was reinstated, allowing for examination of resilience to *in-vitro* stroke-like conditions and BBB repair. **B.** Measurements of tubule permeability were obtained using BI assays before, during, and following *in vitro* strokes using the C10 confocal imager. Representative images are shown for baseline, 5hrs, 8hrs, 72hrs, and 2 weeks after *in-vitro* stroke-like conditions n=5, (right panels), and quantifications of relative permeability (left graph) (n=10 chip cultures/condition/timepoint). Scale= 250 µm **C.** Tight junction morphology was analyzed by ZO-1 staining using the C10 confocal imager. The top panels show examples of intact tight junctions, junctions with zigzags, and junctions with gaps. Representative TJ morphology of anti-miR-23b or controls (left panel) on day 21 following *in vitro* stroke-like conditions (left) and quantifications (right), n=3 chips per condition. Scale = 20µm. **D.** Measurement of transcytosis of anti-miR-23b versus control HBMECs at 5 hours post *in vitro* stroke-like conditions. Shown are images of albumin-594 (red) transcytosis in chip images (left) and quantifications (middle left), and individual images of PLVAP, albumin-594 (middle right), and quantification (far right), n=5. Scale = 12.5µm. **E.** Activation of Wnt/β-catenin post *in vitro* stroke-like conditions was examined at baseline, at sub-acute (3 days) and at chronic time point (21 days) of β-catenin and LEF-1/DAPI, magnified images and quantifications (left panels) and tubule images (right panel), using the C10 confocal imaging plate reader, n=5. Scale = 25µm for cells and 50µm for tubules. Graphs are shown as Mean ± SEM of n=3-10 independent biological replicates, *p <0.05, **p < 0.01, and ***p < 0.001, by two-way ANOVA for multiple comparisons, two-tailed Student’s t-test when analyzing two groups, and multiple unpaired t-tests for each TEER timepoint.

Next, we examined TJ morphology in HBMEC 3D tubules under stroke-like conditions. Although total expression and localization of Claudin-5, Occludin, and ZO-1 were not significantly altered during OGD, structural analysis revealed fewer abnormal ZO-1^+^ TJs with "zigzags" and gaps (green) in anti-miR-23b cultures compared with controls 21 days post-stroke (**Figure 7C**). These data suggest that anti-miR-23b preserves TJ morphological architecture even under hypoxic challenge. Because stroke is known to increase transcytosis^46^, we measured albumin uptake using albumin-Alexa594. anti-miR-23b HBMECs showed significantly reduced albumin uptake 5 hours post-OGD relative to controls (**Figure 7D**). IF analysis also demonstrated decreased levels of PLVAP and intracellular albumin in anti-miR-23b cultures (**Figure 7D**), confirming suppression of transcytosis under ischemic conditions.

To determine whether these are downstream of Wnt/β-catenin activity, we measured β-catenin and LEF-1 protein levels in anti-miR-23b and control cultures before and after OGD. LEF-1 expression was modestly reduced in both groups at 3 days post-stroke. By day 21, anti-miR-23b cultures showed significantly higher expression of both β-catenin and LEF-1 than controls (**Figure 7E**). These findings suggest that anti-miR-23b not only preserves baseline Wnt/β-catenin signaling but also promotes its reactivation during later stages of BBB recovery.

In summary, anti-miR-23b preserves BBB integrity in a 3D stroke model by accelerating barrier recovery and reducing breakdown. It does so by maintaining TJ protein structure, suppressing harmful transcytosis, and bolstering Wnt/β-catenin signaling to restore endothelial barrier function after stroke.

### anti-miR-23b enhances tight junction structure and reduces stroke volume and outcome measures in proof-of-concept experiments using an in vivo ischemic stroke model

Ischemic stroke causes irreversible damage to the infarct core and compromises the surrounding penumbra, a region considered therapeutically salvageable. BBB disruption is an early pathological feature that contributes to the conversion of the penumbra into the infarct core and is associated with poor clinical outcomes^58–60^. Because stroke-induced BBB disruption contributes directly to infarct expansion and cognitive decline, we asked whether anti-miR-23b could confer protection in a clinically relevant *in vivo* model of ischemic stroke. To assess the *in vivo* relevance of our *in vitro* findings, we conducted proof-of-concept studies using the transient middle cerebral artery occlusion (t-MCAO) model to test whether prophylactic treatment of the brain endothelium with anti-miR-23b could reduce BBB dysfunction and stroke severity. This stroke model induces transient cerebral ischemia in the cortex and striatum by inserting an intraluminal suture into the internal carotid artery^46^.

To selectively target brain ECs, we used an adeno-associated viral vector (AAV-BR1) developed by the Körbelin lab^79^, which we engineered to express anti-miR-23b or a control miR. A single intravenous injection (10¹¹ gp/mouse) delivered retro-orbitally resulted in robust transduction efficiency (>75%) in the brain endothelium, as confirmed by co-staining with GFP (from AAV-BR1) and Podocalyxin, a vascular marker (**Figure 8A**). In contrast, astrocytes (GS-positive) and neurons (NeuN-positive) were targeted at low levels (<2%) by the AAV-BR1 viral vector (GFP) in our system (**Figure 8A** and **Figure S6A**), validating brain endothelial cell specificity, as previously shown^79–83^. The transgene cassette used affects the number of non-ECs expressing the AAV-BR1-delivered cargo, which may explain why some studies report neuronal infection^84^. The treatment was well tolerated, with mice maintaining stable body weight and showing no significant systemic immune activation, as confirmed by serum cytokine profiling (IL-1β, IL-6, IFN-γ, TNF-α, IL-8, IL-10, or IL-12) up to 3 months post-injection (data not shown). Consequently, AAV-BR1 represents a promising platform for brain endothelial-specific delivery of anti-miR-23b.

**Figure 8.**
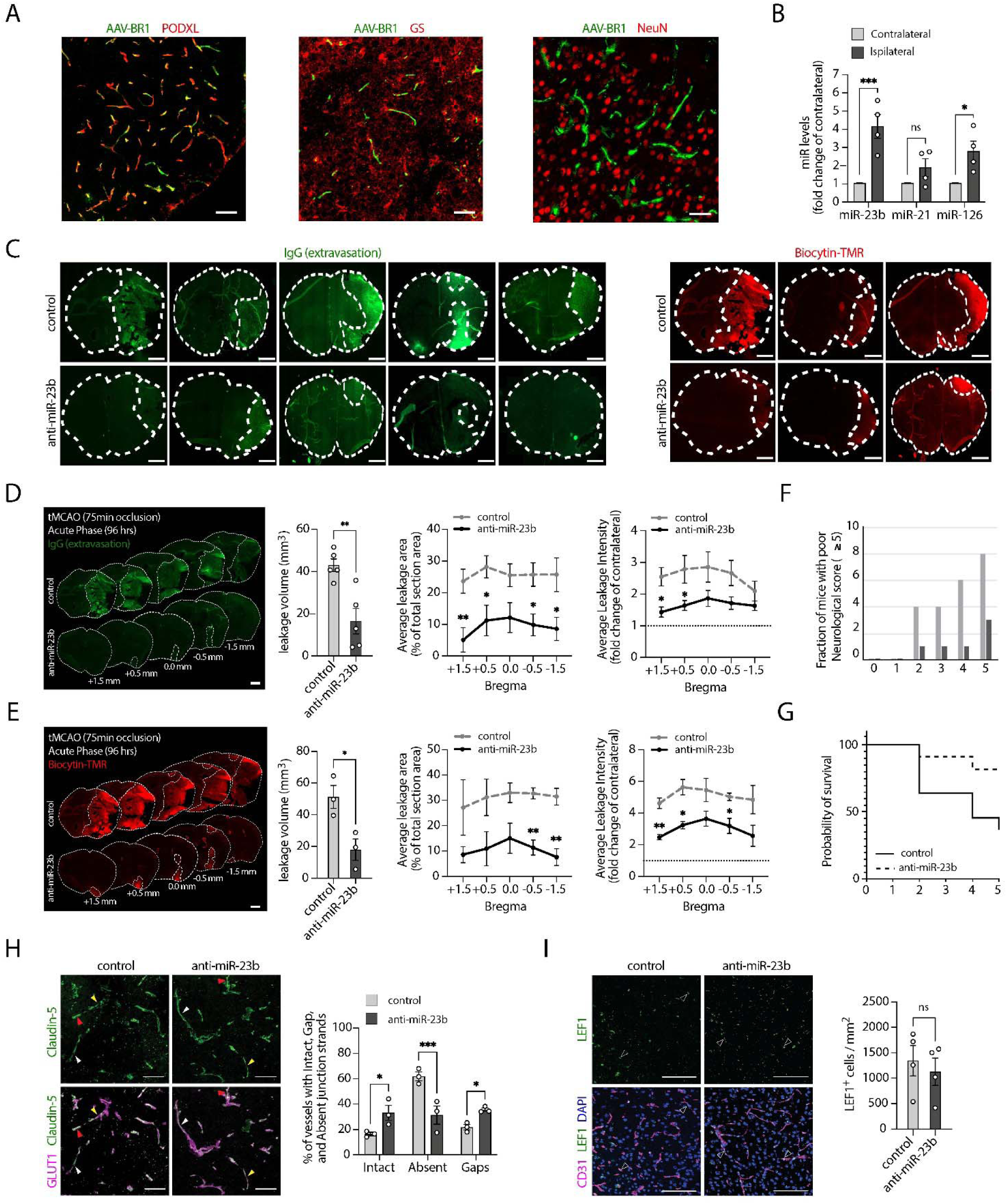
Brain endothelial anti-miR-23b overexpression before stroke reduces acute BBB permeability, enhances tight junction formation and reduces poor outcome measures, following t-MCAO. **A.** Mice were transduced with AAV-BR1 anti-miR-23b or control miR viruses before stroke to target endothelial cells. A representative image of AAV-BR1-control-GFP (green) virus showing expression in brain capillaries co-stained with podocalyxin (PODXL, red vessel marker) left panel, glutamine synthetase (GS, red, astrocytic marker) middle panel, or Neuronal Nuclei (NeuN, red, neuronal marker) right panel, three weeks post-infection, n=3. Scale = 100µm and 50µm (GS and NeuN)**. B.** miR-23b specific qPCR analysis was performed on total brain RNA from PFA sections of control t-MCAO mouse brain, comparing normal vs stroke regions. miR-23b expression levels was normalized to RNU1A, n=3 mice per condition. Two-way ANOVA, bars represent. Bar shows mean ± SEM. **C-D.** 20-week-old male mice were injected with anti-miR-23b-AAV-BR1 or control miR-AAV-BR1 7 days before the t-MCAO surgery (75-minute occlusion). Some mice were sacrificed 4 days after t-MCAO), and BBB damage was examined by analysis of serum IgG extravasation (green). The leakage volume (left panel), average leakage area (middle panel), and average leakage intensity (right panel) for serum IgG extravasation (n=5 mice/group) are shown, Scale = 1mm. Unpaired student t-tests for volume and at each Bregma section for average leakage section and intensity across mice. Paired student t-test for each mouse when comparing contralateral vs ipsilateral sections for average intensity. Bars and XY graphs show Mean ± SEM. **E.** Some mice were exposed to tail-vein injections of a small tracer [biocytin-TMR (red); 890 Da] for 45 minutes before tissue harvest to visualize BBB permeability. Biocytin-TMR leakage volume (left panel), average leakage area (middle panel), and average leakage intensity (right panel, n=3 mice/group), Scale = 1mm. Unpaired student t-tests for volume and at each Bregma section for average leakage section and intensity across mice. Paired student t-test for each mouse when comparing contralateral vs ipsilateral sections for average intensity. Bars and XY graphs show Mean ± SEM. **F.** Neurological scoring was monitored daily. A poor neurological score of> 5 is shown. **G.** Survival monitoring results are depicted in (n=11 mice/group). **H.** Analysis of TJ protein Claudin-5 expression of Glut1 positive brain capillaries were performed comparing control versus anti-miR-23b AAV-BR1 t-MCAO tissue, 3 days post injury. Representative images are shown (left panels) and quantification of tight junction strand structure is shown in the right panel. Intact TJ strands (white arrows), Absent TJ strands (red arrows), and TJ strands with Gaps (yellow arrows) in vascular area of the ipsilateral region of stroke were quantified, n=3 mice per condition. Scale = 100µm. Two-way ANOVA. Bar shows mean ± SEM. **I.** LEF-1 expression was examined in CD31 positive vessels of control vs anti-miR-23b AAV-BR1 treated mice. White arrow represents LEF1+ brain endothelial cells in CD31+ vessels, n=4 mice per genotype. Scale = 100µm. Student t-test. Bar shows Mean ± SEM.

To evaluate therapeutic potential, 3-4-month-old male mice were randomized and injected with AAVBR1-anti-miR-23b or AAVBR1-control-miR one week before 75-minute t-MCAO. Physiological parameters, including cerebral blood flow, temperature, and heart rate, were monitored in a blinded manner, and immune status was assessed using plasma samples (data not shown).

To assess miR-23b expression in the brain, we performed in situ hybridization on healthy and t-MCAO mouse brains. We found that miR-23b mRNA was highly upregulated in a large number of cells in the stroke border region which is a region where angiogenic remodeling of the vasculature is increased together with endothelial Wnt/β-catenin activity (**Figure S6B**). To validate these findings, miR-specific qPCR was performed on total brain tissue from control and t-MCAO subjects. This analysis confirmed significantly elevated levels of miR-23b and miR-126, with modestly increased levels of miR-21 in stroke-affected brains relative to control tissue (**Figure 8B**). In summary, the in situ and qPCR data suggest post-stroke upregulation of miR-23b, consistent with reported increases in plasma miR-23b levels in stroke patients^85^. Consequently, targeting miR-23b with an anti-miR-23b competing RNA may provide therapeutic benefits after a stroke.

A subset of mice was sacrificed 96 hours post-t-MCAO to assess acute outcomes. BBB leakage volume was evaluated by serum IgG extravasation and by biocytin-TMR (870 Da) tracer leakage administered intravenously 45 minutes before euthanasia (**Figure 8C**). Control mice showed extensive BBB leakage in the cortex. In contrast, prophylactic treatment with AAV-BR1-anti-miR-23b significantly reduced the area and intensity of serum IgG and biocytin-TMR leakage (**Figure 8D-E**). Quantitative analysis revealed ∼2-fold reductions in total BBB leakage volume and leakage area across several bregma regions (**Figure 8D & E**) and ∼2-fold reductions in leakage intensity in anti-miR-23b-treated mice compared to controls (**Figure 8C & E**). Importantly, post-t-MCAO barrier integrity analysis (via tracer/IgG extravasation) recapitulated the tracer/stroke volume results from our *in vitro* studies.

Alongside t-MCAO stroke volume analysis, we evaluated neurological scores and survival rates in mice. Compared with controls, the anti-miR-23b group had a lower percentage of animals with severe neurological impairment (score ≥5) and higher survival rates (**Figure 8F**). These findings suggest that anti-miR-23b provides at least some acute neuroprotection (**Figure 8G**).

To further evaluate the *in vivo* relevance of our *in vitro* stroke model, we assessed Claudin-5 expression in GLUT1-positive brain capillaries in tissue from control versus AAV-BR1-anti-miR-23b-treated mice after t-MCAO. anti-miR-23b treatment resulted in a significant increase in intact TJ strands and a corresponding reduction in absent TJ protein strands compared to controls (**Figure 8H**); nonetheless, localized gaps in TJ strands persisted in a subset of the anti-miR-23b group. Post-t-MCAO, we also assessed LEF-1 expression in CD31-positive brain vessels, comparing control and anti-miR-23b-treated mice. No significant difference was observed between the groups (**Figure 8I**). This aligns with our *in vitro* stroke model results, where LEF-1 and β-catenin protein levels showed only modest, non-significant increases at 3 days post *in vitro* stroke, in contrast to day 21 post stroke, when both proteins were significantly upregulated. Despite a limited sample size, we successfully recapitulated key *in vitro* findings in our *in vivo* stroke model, demonstrating that anti-miR-23b modulates BBB permeability (stroke volume) and TJ protein structure. Using these functional metrics, we can evaluate the therapeutic efficacy of anti-miR-23b regimens in maintaining cerebrovascular integrity and facilitating recovery. Additional validation is required, including longitudinal behavioral assessments post-stroke to establish a translational relevance a develop a translational pipeline for anti-miR-23b-mediated restoration and preservation of the BBB.

In summary, our preliminary *in vivo* findings indicate that prophylactic treatment with AAV-BR1-delivered anti-miR-23b effectively mitigates BBB leakage and reinforces tight junction structure. Given the enhanced outcomes in the t-MCAO stroke model, this warrants further investigation of anti-miR-23b as a promising therapeutic strategy for ischemic stroke.

## DISCUSSION

Although acute BBB dysfunction is recognized as one of the earliest and most detrimental events in ischemic stroke^5–7^, a definitive understanding of its mechanisms, timing, and optimal treatment strategies for both protection and repair remain elusive. Our findings establish that targeted silencing of miR-23b in the brain endothelium enhances BBB resilience to injury and facilitates repair through both Wnt/β-catenin-dependent and -independent mechanisms. Inhibiting miR-23b in BECs strengthens the BBB by upregulating critical junction proteins (ZO-1, Claudin-5, Occludin, and VE-Cadherin) and reducing PLVAP-mediated transcellular transport. These protective effects, observed in both healthy and ischemic environments, significantly improve the structural and functional integrity of the BBB. Conversely, overexpression of miR-23b impairs barrier function, confirming its role as a negative regulator of BBB stability.

We developed an unbiased anti-miR screening strategy and identified anti-miR-23b/miR-23b as a conserved anti-miR/miR with potent regulatory activity of endothelial barrier properties, including barrier integrity. Other miRs, including miR-129, miR-221, and miR-191, have also been implicated in regulating vascular responses in stroke pathology, including angiogenesis, highlighting a complex miR regulatory network governing BBB function in neurological diseases^61–63^. However, these miRs were not identified in our anti-miR screening strategy targeting regulators of endothelial barrier integrity. miR-specific qRT-PCR analyses revealed that miR-23a, miR-23b, and miR-27 (which belong to the same miR clusters^41^) are highly expressed in young mouse BECs. However, their levels are reduced in aged BECs, even though they remain highly expressed, suggesting that downregulation of their expression may also be a critical event in promoting the acquisition and maturation of barrier properties of BEC.

Our comprehensive mechanistic study confirms the hypothesis that miR-23b is an endogenously expressed miR that negatively regulates barrier integrity in both mouse and human BECs. miRs can function either by directly binding to and suppressing the translation of specific mRNAs or by targeting mRNAs that influence broader signaling pathways^89,90^. Our multi-pronged approach to assess whether the observed miR/anti-miR effects stem from direct versus indirect regulation demonstrates that miR-23b directly targets VE-Cadherin/CDH5, β-catenin/CTNNB1, and LEF-1 mRNA through imperfect seed interactions, whereas Claudin-5, Occludin, and PLVAP appear to be regulated indirectly through endothelial Wnt/β-catenin activity. Surprisingly, ZO-1/TJP1 was not identified as a direct target in miR eCLIP-seq analysis. However, reporter assays and bioinformatic analyses suggest that miR-23b may bind TJP1 mRNA under certain cellular conditions.

We adopted the unbiased miR-23b eCLIP-seq approach to further examine immediate/early functional outputs elicited by miR-23b or anti-miR-23b in BECs, showing that anti-miR-23b functions as a master regulator of BEC by controlling key signaling pathways (TGF-β, VEGF, Notch, IL-6), in addition to the Wnt/β-catenin pathway. The Wnt/β-catenin, TGF-β, and Notch signaling pathways cooperate to maintain barrier stability and prevent leakiness. In contrast, VEGF and IL-6 signaling generally disrupt the BBB but can also promote beneficial angiogenesis and neuroprotection following injury.

To determine whether miR-23b directly affects Wnt/β-catenin activity, we performed a Wnt-GFP reporter assay. We then used Wnt/β-catenin rescue experiments to evaluate the Wnt-dependent and Wnt-independent contributions to anti-miR-23b barrier function, demonstrating that barrier function is partially Wnt-dependent.

We leveraged multi-omics datasets to critically assess the multi-pronged role of anti-miR-23b in protective barrier function: first, early/intermediate outputs identified by miR-eCLIP-seq analysis showed direct effects on BBB/TJ formation, cell-cell adhesion, transcytosis, and ECM, as well as global effects on Wnt/β-catenin signaling pathways, modulating numerous key mRNAs/proteins (including CTNNB1, LEF-1, WNT7A, FZD4, LRP5, LRP6, APC, TCF3, TCF4, DKD1, WNT5A, ROCK1, DVL2, DVL3, and DAAM1). Additionally, anti-miR-23b regulates the TGF-β and Notch pathways, which integrate with Wnt signaling to maintain barrier stability. In contrast, VEGF and IL-6 pathways, also regulated by anti-miR-23b, typically increase BBB permeability but can also promote protective angiogenesis and neurogenesis. The integrative multi-omics analysis robustly validated that miR-23b is a potent inducer of angiogenesis-related pathways in BECs, while anti-miR-23b effectively inhibits angiogenesis genes and pathways and strengthens a subset of BBB gene signatures. These findings align with previous studies showing that reducing pro-angiogenic pathways including Wnt/β-catenin signaling in BECs is essential to promote and maintain BBB integrity.

Our detailed profiling of miR-23b and its inhibitor in BECs underscores the context- and tissue-specific activities of miR-23b, shedding new light on the seemingly contradictory roles of miR-23 in angiogenic regulation^40,64,65^. Our results indicate that inhibiting miR-23b is beneficial in stroke models, which contrasts with evidence that its induction promotes neuroprotection in Alzheimer’s disease and traumatic brain injury^29,32,66–68^. These disparities likely stem from distinct, cell-type-specific targets of miR-23b, underscoring the need for nuanced, context-dependent approaches to miR-based therapeutics.

Although our analysis focused on anti-miR-23b-mediated inhibition in BECs, potential cross-reactivity with miR-23a, due to their single-nucleotide difference, cannot be ruled out. However, we anticipate minimal impact from inhibiting both variants, given their substantial target overlap. Future studies will confirm this assumption.

Several miRs have already been implicated in regulating stroke outcomes. miR-15a and miR-124 have been associated with ischemic stroke progression and protection, respectively^61,69–71^. miR-129, miR-221, and miR-191 regulate vascular responses during stroke pathology^86–88^. Our data expand on previous reports showing that global, non-specific lentiviral inhibition of miR-23b alleviates ischemic injury by enhancing neuronal survival through the Nrf2 and c-Myc signaling pathways^85,98^. Interestingly, elevated serum levels of miR-23b-3p are observed in patients with ischemic stroke and transient ischemic attack, suggesting a potential role in hypoxia-induced BBB disruption^85^. Yet miR-23b also promotes myelination and neuroprotection^100^, highlighting a complex and paradoxical role for miR-23b across different CNS cell types. Our findings from both the *in vitro* model under stroke-like conditions and the *in vivo* stroke model (t-MCAO) using a brain endothelial-specific approach provide significant insight into the mechanisms by which miR-23b inhibition may confer enhanced BBB stability and possible neuroprotection in ischemic stroke. With miR-23b serving as a negative regulator of BBB integrity, inhibition of miR-23b protects the BBB by enhancing resistance to injury and facilitating repair. It acts by modulating BBB-specific proteins and activating key pathways, such as the Wnt/β-catenin pathway, to promote BBB structural stability and recovery. However, additional validation is required, including longitudinal behavioral assessment following injury to demonstrate translational relevance and to develop the needed translational pipeline for anti-miR-23b-mediated repair and rejuvenation of the BBB.

A major bottleneck to understanding BBB pathology in stroke is the lack of human-relevant, high-fidelity *in vitro* models. Current *in vivo* models, including the t-MCAO model, are valuable but limited by species-specific differences and labor-intensive methods that prevent extensive drug screening^101,102^. Our 3D microfluidic system, bridging simpler 2D assays and complex *in vivo* models, serves as an ideal intermediate platform for studying stroke pathology and screening therapeutic candidates. Our 3D microfluidic BBB model significantly improves on traditional 2D static cultures, providing a robust system for studying mechanisms of ischemic stroke. Our studies demonstrate the value of this 3D BBB culture model as a predictive and mechanistically robust platform, enabling detailed exploration of brain endothelial responses to ischemic injury and facilitating high-throughput screening of next-generation therapeutics. Consistent with our 3D BBB model findings, our *in vivo* study shows that targeting BECs with anti-miR-23b preserves tight junction structure, reduces vascular leakage and stroke volume, and ultimately improves outcomes in ischemic stroke. While anti-miR-23b shows promise as a therapeutic avenue for ischemic stroke, future enhancements should focus on increasing physiological relevance. Integrating a more comprehensive NVU/CNS model that incorporates BBB-supporting cells (pericytes and astrocytes) alongside neurons and microglia will be critical to advancing this therapeutic potential.

The therapeutic potential of anti-miR strategies is illustrated by existing miR-targeted therapies, such as anti-miR-122 (Miravirsen), which has proven safe and effective against Hepatitis C^103–106^. Advanced delivery strategies, such as circular anti-miRs or nanoparticle-based approaches, could enhance therapeutic specificity and efficiency for CNS targeting. There is also significant interest in identifying Wnt/β-catenin agonists that restore homeostatic signaling without overstimulation to promote BBB repair, particularly after stroke. Recently, Wnt7a has been engineered to interact with Grp124/Reck, selectively targeting BEC to promote Wnt/β-catenin signaling and BBB repair in ischemic stroke^17^. Targeting endogenous BEC molecules, such as miR-23b, may offer another safe and effective strategy to achieve this therapeutic goal. Our proof-of-concept studies employing targeted anti-miR-23b delivery via AAV-BR1 specifically improved BBB integrity and outcomes in the t-MCAO model, suggesting a clinical potential. Future research will explore post-stroke therapeutic interventions using anti-miR-23b, including evaluations in middle-aged and aged mice of both sexes. These studies will comprehensively assess neuronal injury, inflammation, immune cell infiltration, and long-term behavioral and cognitive outcomes, thereby significantly advancing translational potential. Ultimately, RNA-based neuroprotection, such as anti-miR-23b, offers superior long-term therapeutic potential compared with conventional pharmacological interventions, which are severely limited by short systemic half-lives. Localized delivery techniques, specifically catheter-based administration during mechanical thrombectomy, could optimize clinical application and significantly improve repair of neurovascular integrity.

In conclusion, given that BBB dysfunction is a critical factor in aging and in neurological conditions, including TBI, MS, Alzheimer’s, and NeuroHIV, our findings represent a significant advance toward therapeutic, small RNA-based gene therapy strategies to restore BBB integrity across multiple CNS pathologies.

## MATERIALS AND METHODS

### Cell Culture

Mouse lung endothelial cells (mLEC; C57-6011, Cell Biologics, Chicago, IL, USA) were maintained in complete mouse EC media (M1168, Cell Biologics, Chicago, IL, USA) and 10% FBS (FB-02, Omega Scientific, Tarzana, CA, USA). Human lung EC (human LEC; #3000, ScienCell, Carlsbad, CA, USA) were maintained on plates coated with 10μg/ml fibronectin (F2006, Sigma-Aldrich, St. Louis, MO, USA) in EC media (1001, ScienCell). Primary human Brain Microvascular Endothelial Cells (HBMEC, ACBRI76, Cell Systems, Kirkland, WA, USA) were cultured on 15μg/ml fibronectin coated T75 flasks (353112, Corning, Corning, NY, USA) in EndoGro-MV Complete Medium (SCME004, Sigma-Aldrich, St. Louis, MO, USA) supplemented with 5ng/mL bFGF (3718-FB-025, R&D Systems, Minneapolis, MN, USA). Human Cerebral Microvascular Endothelial Cell line (hCMEC/D3; SCC066, Sigma-Aldrich, St. Louis, MO, USA) cells were grown on Rat Collagen Type I (C3867-1VL, Sigma-Aldrich, St. Louis, MO, USA) coated T75 flasks in EndoGro-MV complete Medium supplemented with 5ng/mL bFGF. Primary Human Umbilical Cord Vein Endothelial Cells (HUVEC, PCS-100-010, ATCC, Manassas, VA, USA) were cultured on 1% Gelatin (G9136, Sigma-Aldrich, St. Louis, MO, USA) coated T75 flasks in Endothelial Cell Growth Medium MV2 (C-22022, PromoCell, Heidelberg, Germany). HBMECs and HUVECs were used between passages 4 and 8. C57BL/6 Mouse Embryonic Brain Endothelial Cells (C57-6023E, Cell Biologics, Chicago, IL, USA) and Aged Mouse Brain Microvascular Endothelial Cells (A57-6023, Cell Biologics, Chicago, IL, USA) were cultured on 1% Gelatin coated T75 flasks and grown in complete Endothelial Cell Medium (1001, ScienCell, Carlsbad, CA, USA). HEK293T cells (CRL-3216, ATCC) were cultured on 1% gelatin coated flasks in DMEM/F12 medium supplemented with 10% FBS. All cells were cultured at 37°C and 5% CO_2_. The Wnt inhibitor XAV-939 (5uM, 3748, R&D Systems, Minneapolis, MN, USA) or Wnt agonist Wnt3a (100ng, 5036-WN, R&D Systems, Minneapolis, MN, USA) were added to some cultures.

### Cell Transductions and Transfections

#### Stable transduction of miRs

VSV-G-pseudotyped lentiviral particles were made by transfecting 5.3 μg of pMD2-G (12259, Addgene, Watertown, MA, USA), 9.7μg of pCMV-DR8.74 (8455, Addgene, Watertown, MA, USA), and 15μg of pCD510B-1 (miR Control), mZIP, pCD510B-1-miR-23b, or mZIP-anti-miR-23b into 293T cells using Lipofectamine LTX (15338030, ThermoFisher, Waltham, MA, USA). Viral supernatants were concentrated using PEG-it (LV810A-1, System Biosciences, Palo Alto, CA, USA). Cells were transduced with high-titer virus using polybrene (SC-134220, Santa Cruz Biotechnology, Dallas, TX, USA) and spinoculated at 800 × g at 32°C for 30 minutes. HBMEC and hCMEC/D3 transduced cells were selected at 48 hours and maintained with 1 or 3 μg/mL puromycin (J67236.8EQ, ThermoFisher, Waltham, MA, USA), respectively.

### miR mimic transfection

HUVECs were seeded at 300K/well in a 6-well plate and grown until confluent. Transfections were prepared using Lipofectamine™ RNAiMAX Transfection Reagent kit (13778030, ThermoFisher, Waltham, MA, USA) and using both Anti-miR™ miRNA-23b Inhibitor (AM17000, ThermoFisher, Waltham, MA, USA) and Anti-miR™ miRNA Inhibitor Negative Control #1 (AM17010, ThermoFisher, Waltham, MA, USA). The final miRNA concentration added to each well was 25 pmol/well, and the final Lipofectamine™ RNAiMAX added to each well was 7.5μL/well.

### anti-miR lentiviral library screen

mLECs were plated at 10K cells per well in a 96-well plate, incubated with 8μg/mL polybrene (SC-134220, Santa Cruz Biotechnology, Dallas, TX, USA) and transduced by spinoculation (800 x g at 32°C for 30 min) with the miRZipTM anti-miR pooled lentiviral library (MZIPPLVA-1, System Biosciences, Palo Alto, CA, USA) using 10,000 IFUs per well to ensure an MOI<1. Puromycin selection (1μg/mL) was initiated 48 hours after transduction with full media changes every second day. copGFP expression identified cells which had been successfully infected (CopGFP (ppluGFP2) is a bright, green fluorescent protein derived from the copepod *Pontellina plumata*, known for its fast maturation and high stability). Cells were expanded and then single-cell dilution was performed in 96-well format. Single cell clones were expanded and when confluent, cells were transferred to 96-wells allowing for trans-endothelial electrical resistance (TEER) measurements (96W20idf PET plates, Applied BioPhysics). The plates were stabilized with 10 mM L-Cysteine (C6852, Sigma-Aldrich) coated with 100μg/mL Poly-D-lysine (P6407, Sigma-Aldrich), then coated with 10μg/mL fibronectin (F2006, Sigma-Aldrich). For all ECIS experiments, endothelial cells were plated at 50,000 cells/well. TEER was measured at 4000 Hz for 72–96 hrs. For all ECIS experiments, endothelial cells were plated at 50,000 cells/well. anti-miR hits were identified by cloning and sequencing. Next, control miR, miR-23b overexpression and anti-miR-23b (knockdown) high-titer viruses were generated, and human lung endothelial cells (hLECs) were transduced and selected with 10 μg/ml puromycin, after which an ECIS experiment was performed. Finally, HUVEC were transfected with control or anti-miR-23b oligonucleotides, and control or anti-miR-23b HBMEC transduced cells were challenged to TEER analysis.

### Cell proliferation analysis

Cells were plated at low density (6-well plates at 20K cells/well) in triplicate and counted.

### TEER analysis

#### ECIS analysis

The Electrical Cell-Substrate Impedance Sensing system (ECIS ZΘ; Applied Biophysics, Troy, NY, USA) was used to monitor transendothelial electrical resistance (TEER) in real time in 96-well plate format (96W20idf PET plates, Applied Biophysics, Troy, NY, USA). The plates were stabilized with 10 mM L-Cysteine (C6852, Sigma-Aldrich, St. Louis, MO, USA) coated with 100μg/mL Poly-D-lysine (P6407, Sigma-Aldrich, St. Louis, MO, USA), then coated with 10μg/mL fibronectin (F2006, Sigma-Aldrich, St. Louis, MO, USA). All ECIS experiments had lung endothelial cells (LECs) plated at 50,000 cells/well. TEER was measured at 4000 Hz for days to weeks.

### EVOM3 analysis

Parental and miR-modulated HBMECs and HUVECs were both seeded on 15μg/mL Fibronectin or 0.1% Gelatin-coated Transwell® inserts in a 12-well plate (3422, Corning, Glendale, AZ, USA) at 300K cells/well. TEER measurements were performed using an EVOM3 voltohmmeter (World Precision Instruments, Sarasota, FL, USA) at the start of culture and every 24 hours thereafter. Resistance values (Ω) were multiplied by 1.13cm^2^, the surface area of the Transwell® insert, to obtain TEER values (Ω x cm^2^). TEER values of an empty insert were subtracted from each measurement.

### OrganoTEER- 3D TEER

HBSS was added to the middle and bottom channel inlets and outlets (50μL) of the OrganoPlate 3-lane 40/64-chip cultures, and fresh HBMEC-BM was added to the top channel inlets and outlets (50μL). The OrganoTEER device was placed on top of the OrganoPlate, and continuous TEER (Ω x cm^2^) measurements were taken every 8 minutes using the "low TEER" setting. Alternatively, snapshot TEER measurements were performed intermittently, when multiple chip culture plates were analyzed in parallel.

### RNA isolation

Total RNA was isolated from HBMECs (ACBRI76, Cell Systems, Kirkland, WA, USA), Mouse Embryonic Brain ECs (C57-6023E, Cell Biologics, Chicago, IL, USA), and Aged Mouse Brain ECs (A57-6023, Cell Biologics, Chicago, IL, USA) following the manufacturer’s protocol from *mir*Vana™ miRNA Isolation Kit (AM1560, ThermoFisher, Waltham, MA, USA). Alternatively, RNA was isolated from both contralateral and ipsilateral sides of PFA fixed mouse brain tissue by treating tissue with lysis buffer containing 0.5mg/ml proteinase K (4333793, Invitrogen, Waltham, MA, USA) overnight at 60c. The following day, RNA was isolated using *mir*Vana™ miRNA Isolation Kit according to the manufacturer’s instructions. Total RNA was quantified using a NanoPhotometer (IMPLEN, Munich, Germany). Next, cDNA was made using miRCURY LNA RT Kit (339340, Qiagen, Venlo, Netherlands) and qPCR was carried out using the miRCURY LNA SYBR Green PCR Kit (339346, Qiagen, Venlo, Netherlands) and the following miRCURY LNA miRNA PCR Assays from Qiagen: hsa-miR-181d-5p (YP00204789), hsa-miR-149-5p (YP00204321), hsa-miR-24-3p (YP00204260), has-miR-9-5p (YP00204513), has-miR-126-5p (YP00206010), has-miR-21-5p (YP00204230), has-miR-17-5p (YP02119304), hsa-let-7b-3p (YP00205653), hsa-miR-23a-3p (YP00204772), hsa-miR-27b-3p (YP00205915), hsa-miR-23b-3p (YP02119314), hsa-miR-29a-3p (YP00204698), hsa-miR-24-1-5p (YD00610842), hsa-miR-301a-3p (YP00205601), hsa-miR-375-3p (YP00204362), has-miR-151a-3p (YP00204576) and RNU1A1 (YP00203909), on a StepOnePlus™ Real-Time PCR System (4376600, Applied Biosystems, Waltham, MA, USA). All miRCURY LNA miRNA PCR Assays were confirmed to target human and mouse sequences. Each assay was carried out in triplicate, and gene expression for each miR was normalized to the reference gene RNU1A1.

### Western Blotting

Cells were lysed in cold RIPA buffer (89901, ThermoFisher, Waltham, MA, USA) containing protease inhibitor cocktail (PI78410, ThermoFisher, Waltham, MA, USA) to extract protein. Samples containing 1x LDS sample buffer (NP0008, ThermoFisher, Waltham, MA, USA) were boiled at 95 °C for 5 minutes. SDS-PAGE was performed by loading samples on NuPAGE Novex 4-12% Bis–Tris Protein Gels (NP0335, ThermoFisher, Waltham, MA, USA). Methanol-activated PVDF membranes (1704156EDU, Bio-Rad, Hercules, CA, USA) were used to transfer gel samples using the Trans-Blot Turbo Transfer System (1704150, Bio-Rad, Hercules, CA, USA). PVDF membranes were blocked for one hour with 5% non-fat milk (NC9952266, Lab Scientific, Waltham, MA, USA) and incubated overnight at 4°C with the following primary antibodies: ZO-1 (61-7300, Invitrogen, Waltham, MA, USA), Claudin-5 (35-2500, Invitrogen), Occludin (33-1500, Invitrogen, Waltham, MA, USA), GAPDH (MA5-15738, Invitrogen, Waltham, MA, USA), VE-cadherin (2500, Cell Signaling Technology, Danvers, MA, USA), PLVAP (HPA002279, Sigma-Aldrich, St. Louis, MO, USA), LEF1 (2230, Cell Signaling Technology, Danvers, MA, USA), β-catenin (AF1329, R&D Systems, Minneapolis, MN, USA), Cav-1 (3267, Cell Signaling Technology, Danvers, MA, USA). PVDF membranes were washed three times with TBS (1706435, Bio-Rad, Hercules, CA, USA) containing 0.1% Tween (AAJ20605AP, Fisher Scientific, Waltham, MA, USA), then incubated for an hour at room temperature with the following secondary antibodies: Peroxidase AffiniPure™ Donkey Anti-Rabbit IgG (H+L) (711-035-152, Jackson ImmunoResearch, West Grove, PA, USA), Peroxidase AffiniPure™ Donkey Anti-Mouse IgG (H+L) (715-035-150, Jackson ImmunoResearch, West Grove, PA, USA), Goat IgG HRP-conjugated Antibody (HAF109, R&D Systems, Minneapolis, MN, USA). After three additional TBST washes, Pierce ECL Western Blotting Substrate (32106, ThermoFisher, Waltham, MA, USA) and ChemiDoc MP (Bio-Rad, Hercules, CA, USA) gel-documentation system were used for protein expression imaging.

### Immunocytochemistry

HBMECs in the OrganoPlate or on 96-well imaging plates (#655090, Greiner, Kremsmünster, Austria) were fixed with 4% paraformaldehyde in PBS or 100% methanol (-20°C), and permeabilized with 0.2% Triton X-100. Cells were incubated with blocking solution (2% FBS, 2% bovine serum albumin (0332-25G, VWR, Radnor, PA, USA), 0.1% Tween-20) for 30 minutes. Cells were incubated in the following primary antibodies diluted in blocking solution: ZO-1 Monoclonal Antibody (33-9100, Invitrogen, Waltham, MA, USA), Claudin-5 Monoclonal Antibody (35-2500, Invitrogen, Waltham, MA, USA), VE-Cadherin Monoclonal Antibody (2500, Cell Signaling Technology, Danvers, MA, USA), PLVAP (HPA002279, Sigma-Aldrich, St. Louis, MO, USA), LEF1 (2230, Cell Signaling Technology, Danvers, MA, USA), β-catenin (AF1329, R&D Systems, Minneapolis, MN, USA) for 1-2 hours at room temperature, followed by incubation in the following Alexa Fluor-conjugated secondary antibodies: Goat anti-Mouse IgG (H+L) Cross-Adsorbed Secondary Antibody, Alexa Fluor™ 488 (A-11001, Invitrogen, Waltham, MA, USA), Goat anti-Mouse IgG (H+L) Cross-Adsorbed Secondary Antibody, Alexa Fluor™ 594 (A-11005, Invitrogen, Waltham, MA, USA), Donkey anti-Goat IgG (H+L) Cross-Adsorbed Secondary Antibody, Alexa Fluor™ 647 (A-21447, Invitrogen, Waltham, MA, USA), Donkey anti-Mouse IgG (H+L) Highly Cross-Adsorbed Secondary Antibody, Alexa Fluor™ 555 (A-31570, Invitrogen, Waltham, MA, USA), Donkey anti-Goat IgG (H+L) Cross-Adsorbed Secondary Antibody, Alexa Fluor™ 594 (A-11058, Invitrogen, Waltham, MA, USA), Donkey anti-Rabbit IgG (H+L) Highly Cross-Adsorbed Secondary Antibody, Alexa Fluor™ 488 (A-21206, Invitrogen, Waltham, MA, USA) for 30 minutes at room temperature. Nuclei were stained with DAPI (62248, ThermoFisher, Waltham, MA, USA). Cells were imaged with Cytation C10 Confocal Imaging Plate Reader using 20x objective or Leica DMI8 SP8 Confocal Microscope (Leica, Wetzlar, Germany) using 40x or 63x oil-immersion objectives. Images are the max projection of 10-micron Z-stacks with 1 micron Z-step per image.

### OrganoPlate HBMEC culture

All experiments used the Mimetas OrganoPlate 3-lane platform (4004-400B, MIMETAS, Leiden, Netherlands). ECM gel was prepared on ice using 100mM HEPES (B35110, R&D Systems, Minneapolis, MN, USA), 37 g/L NaHCO_3_ (S5761, Sigma-Aldrich, St. Louis, MO, USA), 5mg/mL Human Collagen Type IV (C5533-5MG, Sigma-Aldrich, St. Louis, MO, USA), and 5mg/mL Cultrex 3-D Culture Matrix Rat Collagen I (3447-020-01, Sigma-Aldrich, St. Louis, MO, USA) in a 1:1:1.6:6.4 ratio. Then, 2μL of ECM gel was dispensed into the middle channel inlet and incubated for 15 min at 37°C. After plate incubation, 50μL of PBS was added to the middle inlet to prevent the gel from drying. Next, 40μL of coating solution containing 50μg/mL collagen type IV and 30μg/mL fibronectin in PBS was used to coat the top channel of every chip, and the plate was left to incubate at 37°C for 1-3 hours. After the coating was aspirated, the top channels were washed with 50μL of PBS. PBS in the top channel inlet was replaced with 50μL of OrganoMedium HBMEC-BM (MI-OM-HBBM, Leiden, Netherlands). An HBMEC cell suspension of 1.5 x 10^6^ cells/mL was prepared in OrganoMedium HBMEC-BM, and 2μL of cells were seeded in each top channel outlet. The OrganoPlate was incubated statically, on its side, for 3 hours to allow cells to adhere to ECM, after which 50μL of OrganoMedium HBMEC-BM was added to the top channel outlets and the OrganoPlate was then placed on OrganoFlow L perfusion rocker (MI-OFPR-L, MIMETAS, Leiden, Netherlands) at a 7° inclination, 8-min interval. Medium was changed every 2-3 days by replacing the top channel OrganoMedium HBMEC-BM in the top channel inlets and outlets with fresh OrganoMedium HBMEC-BM.

## 3D *in vitro* Stroke Assay

Stroke was modeled using a three-fold approach (hypoglycemia, hypoxia, and flow stasis) for 6 hours. First, hypoglycemic conditions were achieved by replacing OrganoMedium HBMEC-BM culture media with deoxygenated Glucose Free Endothelial Cell Medium (1001-GF, ScienCell, Carlsbad, CA, USA). Hypoxic conditions were achieved by placing the OrganoPlate in CellXpert^®^ C170i (Eppendorf, Hamburg, Germany) cell culture incubator set at 1% O_2_. Flow stasis was achieved by removing the OrganoPlate from the OrganoFlow rocker and placing it flat in the incubator. HBMEC barrier integrity was monitored throughout the study by OrganoTEER.

### Transcytosis Assay

Cells grown in 2D monolayers were seeded at 50K cells per well (96-well plate) and cultured until confluent. First, cells were incubated with 50ug/mL Labeled Bovine Serum Albumin CF®594 Dye Conjugate (20290, Biotium, Fremont, CA, USA) at 37°C for 1 hour. For 3D cultures, media containing 50ug/mL Labeled BSA CF®594 Dye Conjugate was added to the top channel inlets and outlets of the OrganoPlate and was allowed to rock for 1 hour. Following incubation, cells were washed of excess BSA-594 and were analyzed for CF®594 Mean Fluorescence Intensity (MFI) using Cytation C10 confocal imager or fixed with 4% PFA and co-stained with an immunofluorescence antibody.

### Tracer Barrier Integrity Assay

At least 12 hours after seeding the 3D OrganoPlate, 0.5μg/mL FITC-Dextran 150kDa (46946, Sigma-Aldrich, St. Louis, MO, USA) in HBSS was introduced into the top channel inlets and outlets (40μL and 30μL). Middle and bottom channel inlets and outlets were filled with 20 μL HBSS. The OrganoPlate was then placed flat for 15 minutes to allow tracer medium to perfuse through the top channel. Fluorescence intensity was measured via an EVOS microscope or Cytation C10 confocal plate reader. Analysis was performed by normalizing the average intensity of the middle and bottom channels over the intensity of the top channel, which represents leakage from the vascular (top) channel into the brain (middle and bottom) channels.

### 3’UTR Luciferase Reporter Assay

TJP1 SC214630, CLDN5 SC207859, OCLN SC208413, CDH5 SC215693, CTNNB1 SC212476, LEF1 SC213363 and control PS100062 3’UTR luciferase containing plasmids was employed in reporter assays. HEK293T cells were co-transfected with 0.8µg of either luciferase reporter plasmids and 10nM miR-23b mimic (MC12931, Invitrogen, Waltham, MA, USA) or control miR mimic (AM17010, ThermoFisher, Waltham, MA, USA) using FuGENE (E5911, Promega, Madison, WI, USA) transfection reagent according to manufacturer instructions. 24 hours after transfection, cells were lysed using Pierce Firefly Luciferase Glow Assay Kit (16176, Invitrogen, Waltham, MA, USA) according to the manufacturer’s instructions. Luminescence was detected by Cytation C10 plate reader.

### Wnt-GFP Reporter Assay in HEK293T cells

HEK293T cells were seeded at 150K cells/well in 12 well dishes the day before transfection. 100 ng of M38 TOP-dGFP (17114, Addgene, Watertown, MA, USA). Wnt reporter vector was transfected into each well along with 10nM of negative control miR or 10nM miR-23b (MC12931, Invitrogen, Waltham, MA, USA) oligonucleotides with or without 100ng/mL of Wnt3a (5036-WN-010, R&D Systems, Minneapolis, MN, USA) using Lipofectamine 3000 (Invitrogen, Waltham, MA, USA). Plates were imaged after 24 hours and GFP intensity per cell was quantified using the Cytation C10 confocal plate reader. The results represent triplicate determination of a single experiment that is representative a total of four similar experiments.

### miR-eCLIP Library Preparation and Analysis Methods

For miR-eCLIP experiments, the standard eCLIP protocol^107^ was modified to enable chimeric ligation of miRNA and mRNA^67^. Studies were performed by Eclipse Bioinnovations Inc. (www.eclipsebio.com, San Diego, CA, USA). Primary HBMECs were transduced with miR control, miR-23b or anti-miR-23b as described above and the functional phenotype on TEER was validated. Cells were UV crosslinked at 400mJoules/cm2 with 254 nm radiation and stored until use at -80°C. Crosslinked cell pellets were then lysed with 750μL of eCLIP lysis mix and sonicated (QSonica, Q800R2) for 5 minutes, 30 seconds on / 30 seconds off with an energy setting of 75% amplitude, followed by digestion with RNase-I (Ambion, Waltham, MA, USA). A primary mouse monoclonal AGO2/EIF2C2 antibody (sc-53521, Santa Cruz Biotechnology, Dallas, TX, USA; H00027161-M01, Novus Biologicals, Centennial, CO, USA) was incubated for 1 hour with magnetic beads pre-coupled to the secondary antibody (M-280 Sheep Anti-Mouse IgG Dynabeads, 11202D ThermoFisher, Waltham, MA, USA) and added to the homogenized lysate for overnight immunoprecipitation at 4°C. Following overnight IP, 2% of the sample was taken as the paired size-matched input with the remainder magnetically separated and washed with eCLIP high stringency wash buffers. Chimeric ligation was then performed on-bead at room temperature for 1 hour with T4 RNA ligase (NEB). IP samples were then dephosphorylated with alkaline phosphatase (FastAP, ThermoFisher, Waltham, MA, USA) and T4 PNK (NEB0) and an RNA adaptor was ligated to the 3’end. 83% of IP samples were used for miR23-capture libraries using antisense probes to miR23. RNA adapter ligation, reverse transcription, DNA adapter ligation, and PCR amplification were performed as previously described. After sequencing, samples were processed with Eclipsebio’s proprietary analysis pipeline (v1). UMIs were pruned from read sequences using umi_tools (v1.1.1). Next, 3’ adapters were trimmed from reads using cutadapt (v3.2). Reads were then mapped to a custom database of repetitive elements and rRNA sequences. All non-repeat mapped reads were mapped to the genome UCSC version GRCh38/hg38 using STAR (v2.7.7a). PCR duplicates were removed using umi_tools (v1.1.1). AGO2 eCLIP peaks were identified within eCLIP samples using the peak caller CLIPper (v2.0.1). For each peak, IP versus input fold enrichments and p-values were calculated. miRNAs from miRBase (v22.1) were "reverse mapped" to any reads that did not map to repetitive elements or the genome using bowtie (v1.2.3). The miRNA portion of each read was then trimmed, and the remainder of the read was mapped to the genome using STAR (v2.7.7a). PCR duplicates were resolved using umi_tools (v1.1.1), and miRNA target clusters were identified using CLIPper (v2.0.1). Each cluster was annotated with the names of miRNAs responsible for that target. Peaks were annotated using transcript information from GENCODE Release 35 (GRCh38.p13) with the following priority hierarchy to define the final annotation of overlapping features: protein coding transcript (CDS, UTRs, intron), followed by non-coding transcripts (exon, intron).

### Bulk RNA Sequencing, Alignment, and Analysis

Total RNA was isolated from miR control, miR-23b and anti-miR-23b overexpressing HBMEC, following the manufacturer’s protocol from *mir*Vana™ miRNA Isolation Kit (AM1560, ThermoFisher, Waltham, MA, USA). RNA samples underwent quality assessment using a NanoDrop spectrophotometers and RNA samples were submitted for RNA sequencing at Azenta (Burlington, MA, USA. Raw paired-end RNA-sequencing reads (FASTQ) were quantified using kallisto (v0.52.0). Transcript-level abundances were summarized to gene-level counts using tximport with a transcript to gene mapping derived from the Homo sapiens GRCh38.105 annotation GTF using txdbmaker. The DESeq2^108^ (v1.50.2) framework was used on RStudio (v4.5.2) for normalization, dispersion estimation, and fitting of negative binomial generalized linear models. Comparisons between miR-23b-overexpressing, anti-miR-23b-overexpressing and control samples were evaluated, and genes were ranked by Benjamini-Hochberg adjusted p-values. Variance-stabilizing transformation (VST) was applied for exploratory analyses, including principal component analysis. Differential expression analysis was performed using the edgeR package, considering Padj <0.05 and a log2-fold change <−0.25 or > 0.25. Adjusted p-values were calculated using the Benjamini-Hochberg procedure implemented in edgeR. Principal component analysis was performed, and Pearson correlation values were computed using the resulting Log2 counts per million values. CPA, Volcano, GSEA plots and Heatmaps were generated with the heatmap package^109^ using min-max scaling on normalized counts. Significance thresholds were set at adjusted p-value < 0.05 and log2 fold change > 0.6.

### Proteomics

#### Chemicals and reagents

Solvents, including water, employed for proteomics sample preparation and chromatography were of Optima LC/MS grade (ThermoFisher, Waltham, MA, USA). Formic acid (A11710X1-AMP, ThermoFisher, Waltham, MA, USA) and trifluoroacetic acid (TFA, 28904, ThermoFisher, Waltham, MA, USA) were obtained in sealed 1 mL ampules. All additional reagents were of analytical grade unless otherwise specified.

### Sample preparation and mass spectrometry

Confluent cells were harvested, washed with PBS buffer and lysed (lysis buffer: 5% SDS, 50mM Triethylammonium bicarbonate). The lysate was heated at 60°C for 30 min on a block heater. Proteins were processed for proteomics as described previously^110^. Following acidification to pH ∼1 with 12% phosphoric acid, the samples were transferred to S-Trap micro columns (P/N C02-micro-80, Protifi, Fairport, NY,). On-column digestion was performed overnight using trypsin and the resulting peptides were eluted with ammonium bicarbonate. The peptides were lyophilized and re-suspended in 3% acetonitrile, 0.1% formic acid. Final peptide concentrations were determined with a NanoDrop Spectrophotometer (ND-ONE-W, ThermoFisher, Waltham, MA, USA). Peptides were analyzed using a Vanquish Neo liquid chromatograph coupled to an Orbitrap Exploris 240 mass spectrometer (ThermoFisher, Waltham, MA, USA). Chromatographic separation was performed on an EASY-Spray PepMap C18 75 µm ID x 50 cm column at a flow rate of 300 nL/min with an acetonitrile/formic acid gradient. The mass spectrometer was operated in data dependent (DDA) mode for spectral library construction and data independent acquisition (DIA) mode for quantitative protein analysis. Ion source conditions were maintained with a capillary voltage of 3400 V, source temperature of 280°C, and an S-Lens RF level of 70%. DDA settings included a cycle time of 1.2 sec, dynamic exclusion for 40 sec, and resolution of 120,000 for MS and 30,000 for MS/MS. In DIA mode, MS full scans were acquired at 12,000 resolution followed by MS/MS scans at 30,000 resolution. DIA settings included a precursor mass range (m/z) of 375-1275, with 60 isolation windows of 5 m/z each, with an overlap of 1 m/z.

### Mass spectrometry data analysis

Raw mass spectrometry data were processed using PEAKS Studio v. 13.0 (Bioinformatics Solutions, Inc., Waterloo, ON, Canada)^111^. Sequences were matched against the Human UniProt reviewed database (Release 2026_01, including isoforms). A spectral library was constructed from DDA files to guide the annotation and quantification of DIA data. Search parameters included carbamidomethylation of cysteine as a fixed modification and oxidation of methionine and protein N-terminal acetylation as variable modifications. Mass tolerances were set at 10 ppm for precursor ions and 0.02 Da for fragment ions. Protein quantification was performed using the MaxLFQ algorithm in PEAKS 13.0, which calculates abundance ratios based on median peptide ratios to minimize the impact of outliers and missing values^112^. Normalization was conducted within PEAKS Studio based on Total Ion Current (TIC)^113,114^. PEAKS used the ANOVA method to calculate the significance and generate a global p-value for each protein. All mass spectrometry raw data have been deposited in an international public repository (MassIVE at https://massive.ucsd.edu) under the accession number MSV000101239.

### Bioinformatics and statistical analysis

Subsequent statistical analyses were conducted in RStudio (v.4.2.3) using the *rstatix* package (v. 0.7.2). To prepare the data for pairwise group mean comparisons, protein abundance values were log10-transformed and then analysed via the Games-Howell post-hoc method. This procedure establishes confidence intervals and assesses significance while inherently adjusting for multiple testing, which eliminated the requirement for additional p-value corrections^115^.

Quantitative proteomics data were analyzed using DEP (v 1.26.0) on R (v 4.4.1.). Proteins were filtered to retain entries quantified across all samples (filter_missval, threshold = 0). Intensities were normalized using variance stabilizing normalization (normalize_vsn) based on the vsn method. Missing values were imputed using a left-censored approach with the impute() function (method = "MinProb", q = 0.01), which replaces missing intensities with values drawn from a distribution representing low abundance proteins. Differential protein expression was assessed using test_diff across anti-miR23b, miR23b, and control conditions. Significant proteins were defined using add_rejections with thresholds of adjusted p-value < 0.05 and log2 fold change > 0.6.

### AAV-BR1 anti-miR-23b or control miR Administration

To target the cerebral endothelium, we generated AAV-BR1 anti-miR-23b and miR control virus (AAV2-BR1 plasmids from Dr. Körbelin’s lab^79^. Initial dosing and safety analysis showed that a single dose (i.v.), (10x^11^ gp/mouse) of AAVBR1 virus encoding anti-miR-23b-GFP (or -mCherry, -Luciferase) was effective (>70% transduction rate) when introduced into mice by retro-orbital injections and analyzed after 3 weeks, demonstrating specific expression in brain endothelium, when stained using podocalyxin, a HBMEC marker and AAV-BR1-GFP. As reported, the AAV-BR1 virus remained expressed for a prolonged period (>600 days). Mice were active and maintained a healthy weight following viral administration. Analysis of the peripheral immune response by cytokine profiling of plasma (ProcartaPlex 36-plex arrays, EPX360-26092-901, Invitrogen, Waltham, MA, USA showed no significant changes in expression of various cytokines (IL-1β, IL-6, IFN-γ, TNF-α, IL-8, IL-10, or IL-12) after 3 months, supporting the notion that the virus is well-tolerated.

### Transient Middle Cerebral Artery Occlusion (t-MCAO)

12 to 14-week-old male mice weighing 25-30g were used. Briefly, mice were anesthetized with 1-2% isoflurane in 100% oxygen. After the surgical exposure, the right external and common carotid arteries were permanently ligated. A reusable monofilament (70SPRe2045, Doccol Inc., Sharon, MA, USA) was introduced into the right common carotid artery and advanced 9–10 mm beyond the carotid bifurcation into the right internal carotid artery. The core body temperature was checked by rectal probe and maintained at 36.5 ± 0.5°C by a heating pad and lamp throughout the surgery. A laser Doppler flowmeter measured the cerebral blood flow pre-occlusion, at beginning of occlusion and post-occlusion, with at least an 80% reduction compared to presurgical baseline values achieved after the insertion of the filament. Specifically, mean % occlusion from mice used was 86.23%. The flowmeter probe was placed 2mm posterior and 5-6 mm lateral to the bregma on the skull. The reperfusion was achieved by withdrawing the filament 75 min after the insertion. Following closure of the surgical wound, mice were injected subcutaneously with 2.5 mg/kg flunixin, recovered on a warming pad, and then returned to their cages with free access to water and food. 30 minutes prior to euthanization, mice received a tail vein injection of 1% Biocytin-TMR in PBS (T12921, ThermoFisher, Waltham, MA, USA). Tissue was obtained at 96 hrs. post-t-MCAO for some animals.

### *In situ* hybridization

In situ hybridization was performed on fresh frozen brain mouse sections at 72 hours post-t-MCAO as described^116^ using miR-23b (mmu-miR-23b-3p miRCURY LNA miRNA Detection probe (YD00611001-BCG, Qiagen) or scramble-miR probe (scramble miR miRCURY LNA miRNA Detection (YD00699004-BCG, Qiagen). The hybridization of the antisense probes on the brain tissue and washings were performed at 50C. The images were acquired on a Zeiss M2 Axioimager.

### Immunohistochemistry

Animals were euthanized with isoflurane and perfused first with PBS, then with 4% PFA in PBS. Brains were fixed in 4% PFA for 24 hours, after which tissue was cryoprotected in 30% sucrose for 24-48 hrs., embedded in Tissue-Tek OCT and frozen for 24 hours at -80 °C. Brains were sectioned in 12 µm-thick coronal slices spanning all bregma regions of interest using a Leica cryostat. Tissues underwent a series of ethanol washes before blocking with PBST (PBS with 0.1% Triton-X) containing 10% BSA. The following primary antibodies were diluted in PBST containing 1% BSA and added to tissue overnight at 4C: anti-NeuN (ab177487, Abcam, Waltham, MA, USA), anti-Podocalyxin (AF1556, R&D Systems, Minneapolis, MN, USA), anti-Glutamine synthetase (ab73593, Abcam, Waltham, MA, USA), anti-LEF1 (2230, Cell Signaling Technology, Danvers, MA, USA), anti-GLUT1 (07-1401, Sigma-Aldrich, St. Louis, MO, USA), anti-CD31 (M0823, Agilent Dako, Santa Clara, CA, USA). For Claudin-5 (Cldn5) Immunostaining, tissue underwent antigen retrieval using 1x citrate buffer (100C for 35 minutes) prior to blocking and incubating with anti-Claudin-5 (35-2500, ThermoFisher Waltham, MA, USA) primary antibody overnight. The next day, Primary antibody signal was enhanced using the following fluorescence-conjugated secondary antibodies: Donkey anti-Goat IgG (H+L) Cross-Adsorbed Secondary Antibody, Alexa Fluor™ 647 (A-21447, Invitrogen, Waltham, MA, USA), Donkey anti-Rabbit IgG (H+L) Highly Cross-Adsorbed Secondary Antibody, Alexa Fluor™ 488 (A-21206, Invitrogen, Waltham, MA, USA) for 2 hours at room temperature. To assess leakage in brain sections, Streptavidin-Alexa Fluor 594 (S11227, Invitrogen, Waltham, MA, USA)was used to amplify the Biocytin-TMR signal, and Goat anti-Mouse IgG (H+L) Cross-Adsorbed Secondary Antibody, Alexa Fluor™ 488 (A-11001, Invitrogen, Waltham, MA, USA) was used to visualize serum IgG in brain sections. Leakage sections were imaged with an upright Zeiss Axioimager. Sections observing vasculature were imaged using a C10 confocal plate reader or a Zeiss LSM700 confocal microscope.

### BBB leakage Image Analysis

Using FIJI, uniformly threshold bregma section images were used to measure the area of fluorescence of each Biocytin-TMR and IgG signal that exceeded the threshold. Fluorescent areas were then normalized to the total area of each bregma section. Biocytin-TMR and IgG percent leakage area were plotted across specific bregma coordinates (+1.5, +0.5, 0.0, -0.5, -1.5). The volume of Biocytin-TMR and IgG leakage in each mouse was calculated by equating leakage volume to that of a truncated cone utilizing the cross-sectional areas and distance between sections. Intensity of each leakage was determined by sampling 10 identical regions in contralateral and ipsilateral cortex of each bregma section and calculating the ratio between the ipsilateral and the contralateral.

### Tight junction morphology

To quantify junctional abnormalities, 8 images/animal were acquired from the ischemic core at bregma -0.5. The percentage of vessel segments with either intact TJ strands or TJ strands with gaps or absent junctional strands over the total number of vessel segments was calculated per image in ImageJ. A “gap” is defined as a continuous GLUT1-positive vessel segment in which Cldn5 intensity fell below a specified threshold while GLUT1 stayed intact. An “absent” junction can be defined as Cldn5 signal that does not resemble a strand-like morphology, but instead an intracellular-like morphology when overlayed with GLUT1-positive vessel segment. An “intact” junction is defined as Cldn5 signal that resembles a strand-like morphology and covers the entirety of a GLUT-1 positive vessel segment.

### Statistics

Data was analyzed using GraphPad Prism (version 11). Individual wells, or chips were considered biological replicates. All key experiments were repeated independently, and detailed information about sample size, error bars, and the number of experiments is provided in each figure legend, while p-values are provided in the figure. The Shapiro-Wilk test was used to assess the normality of the data. For normally distributed data, two-way ANOVA was used to compare the effects and interactions of either miR-23b overexpressing or anti-miR-23b independent variables. A Student’s t-test with Welch Correction was used for comparisons between two groups. For datasets not distributed normally, non-parametric Mann-Whitney U test was used for comparisons between two groups. For *in vivo* mouse analyses, paired t-tests were used to compare stroke versus contralateral regions within the same mouse. Unpaired t-tests were used to compare across different conditioned mice. Outliers were identified using the interquartile range (IQR) method. Data are displayed as mean ± S.E.M. of technical or independent biological replicates (n) as indicated. *p< 0.05, **p< 0.01, ***p < 0.001, or ****p< 0.0001.

## ACKNOWLEDGEMENT

The work of I.M. Pedersen is supported by the National Institutes of Health and DoD (grants R01 NS107344, R61R33 HL159949, HL159949-S1 and TP425-24-1-1046). L.M Brown (5R01AG067501-05, 5R01AG071868-03, 5R01AG073360-04, 5R01DK124465-05, and 1R01AG076949-01/RFMH158825). This is manuscript #1083 from Scintillon Research Institute. This is manuscript #1083 from Scintillon Research Institute.

## AUTHOR CONTRIBUTIONS

V. Martinez designed and performed most of the experiments and analyzed data. R. Presby, J. Krog, I. Daugaard, L. Das and D. Jamoul performed experiments, and P. Gaur, U. Akcan, D.S. Johnson and S. Nakanishi analyzed data. L.M. Brown and V. Menon advised on multi-omics analysis, J. Körbelin provided AAVBR1 plasmids, V. Martinez, R. Presby and S. Nakanishi contributed to writing the methods sections and generating the figures. I.M. Pedersen designed all experiments, analyzed data, and wrote the paper with input from D. Agalliu and B. Lawson.

