## Supplemental Figures for "Modulation of miR-23b Wnt/β-catenin Axis Strengthens Endothelial Barrier Properties"

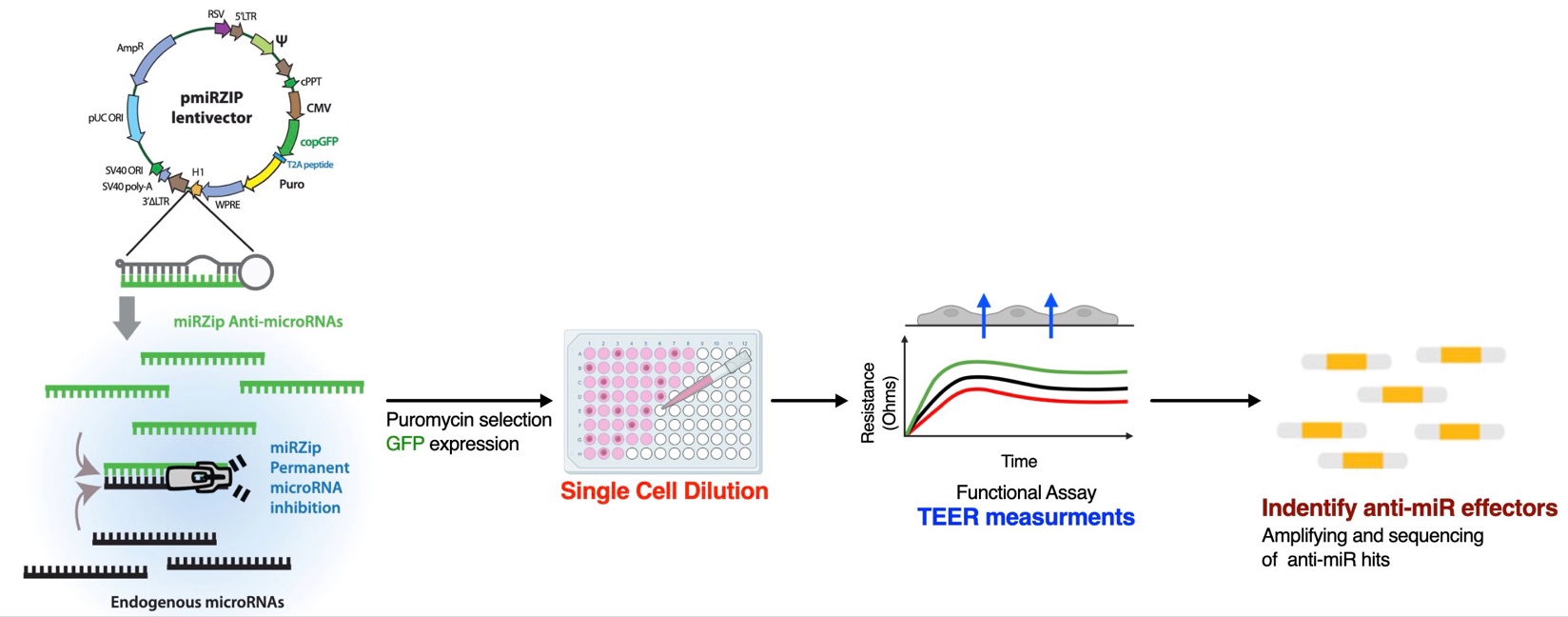
**Supplemental Figures**

**Figure S1.** Schematic diagram of the anti-miR library screen to identify anti-miRs/miRs that regulate cell-cell contact and barrier properties in endothelial cells. Approved modification from cartoon on System Biosciences’ website.

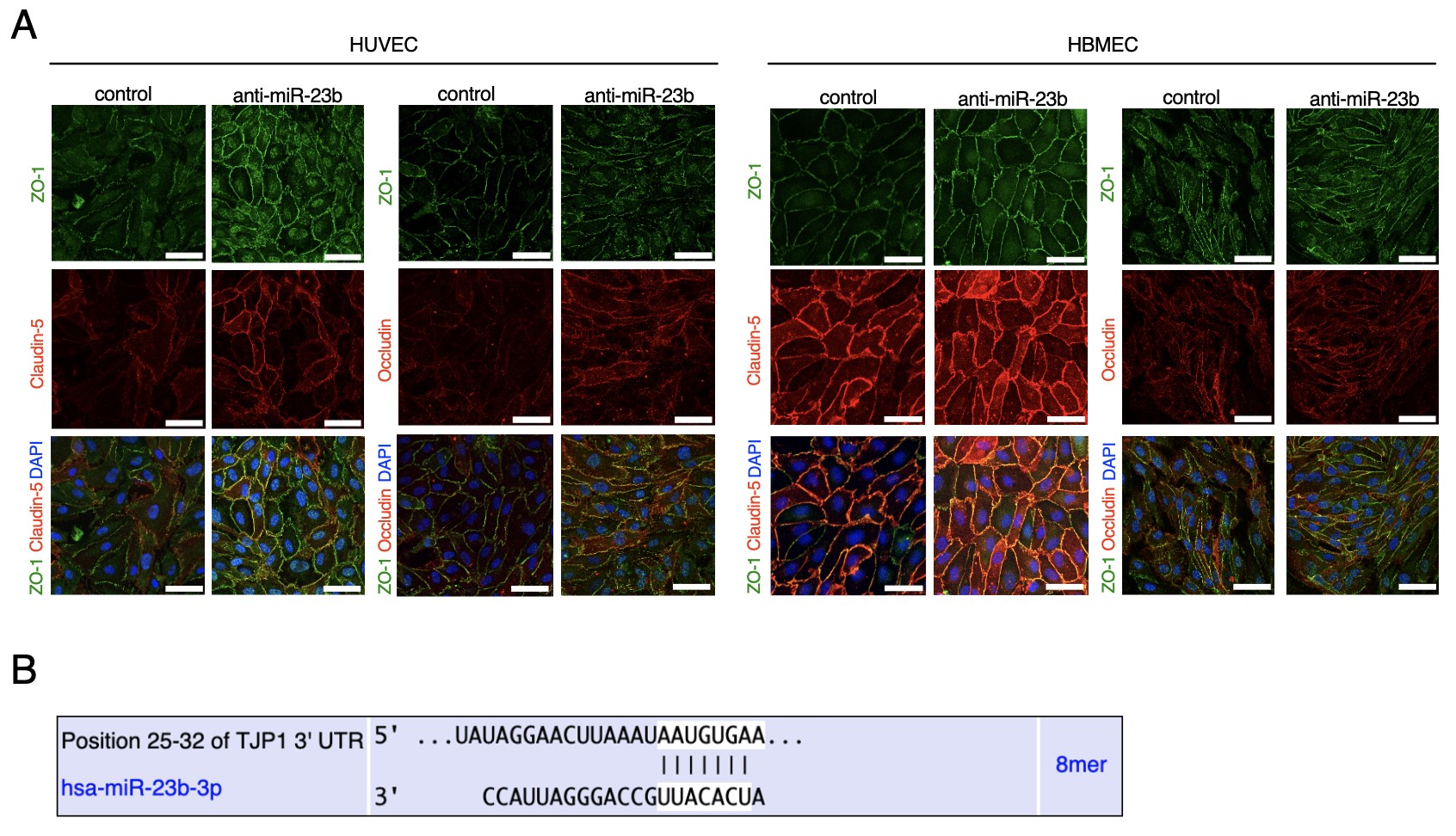

**Figure S2. anti-miR-23b enhances tight junction proteins in ECs – single TJP images, and predicted miR-23b binding site in TJP1/ZO-1 mRNA. A.** Single tight junction protein images (ZO-1, Claudin-5 and Occludin) in anti-miR-23b and control miR modulated HUVEC and HBMEC. Scale = 50µm. **B.** Schematic diagram of predicted miR-23b seed binding site in TJP1/ZO-1 mRNA, from TargetScanHuman 8.0.

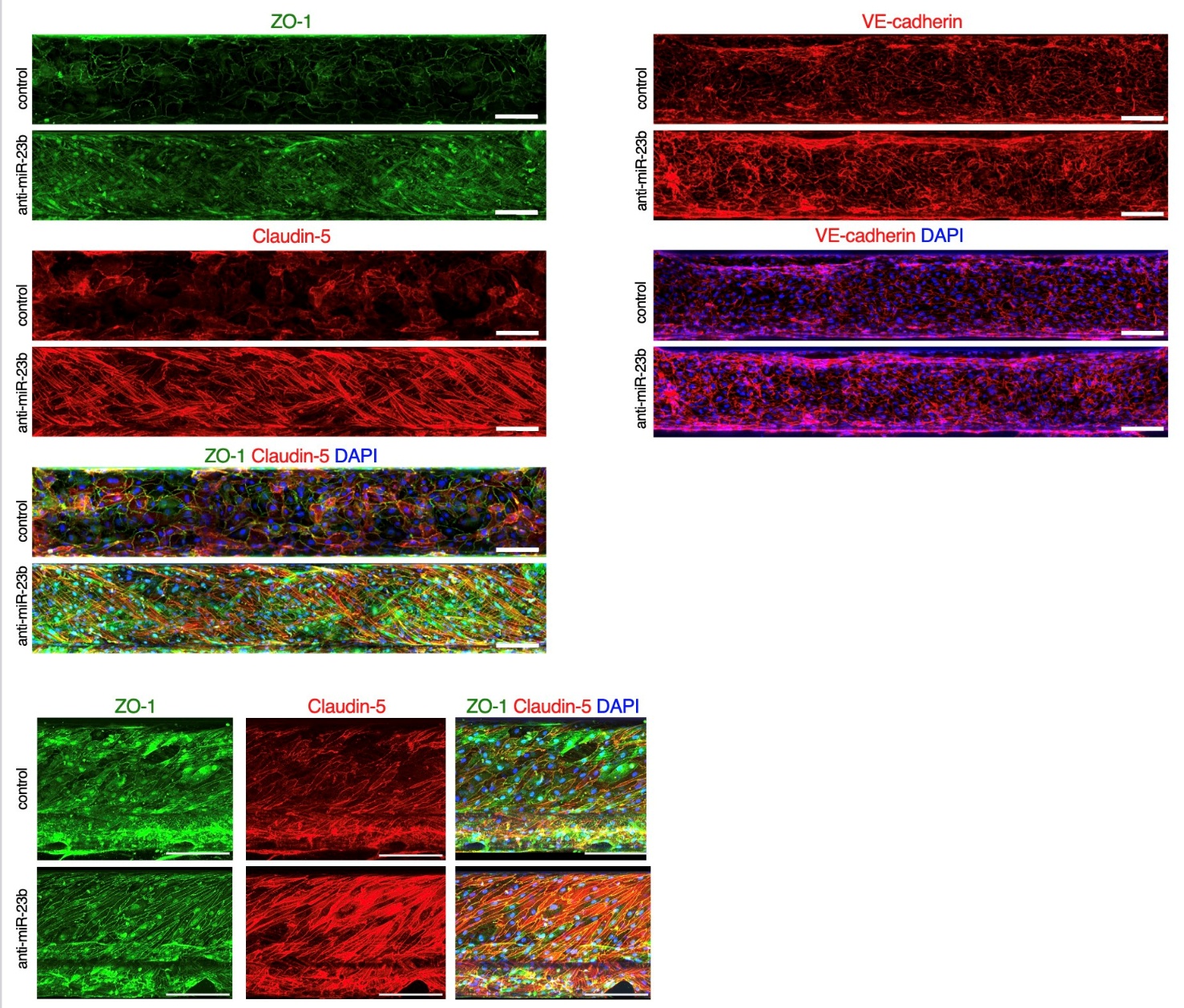

**Figure S3. anti-miR-23b enhances tight junction proteins in 3D tubules – single TJP images.**

Single TJ protein images (ZO-1, Claudin-5 and Occludin) in anti-miR-23b and control miR modulated HBMEC 3D tubules. Scale = 250µm.

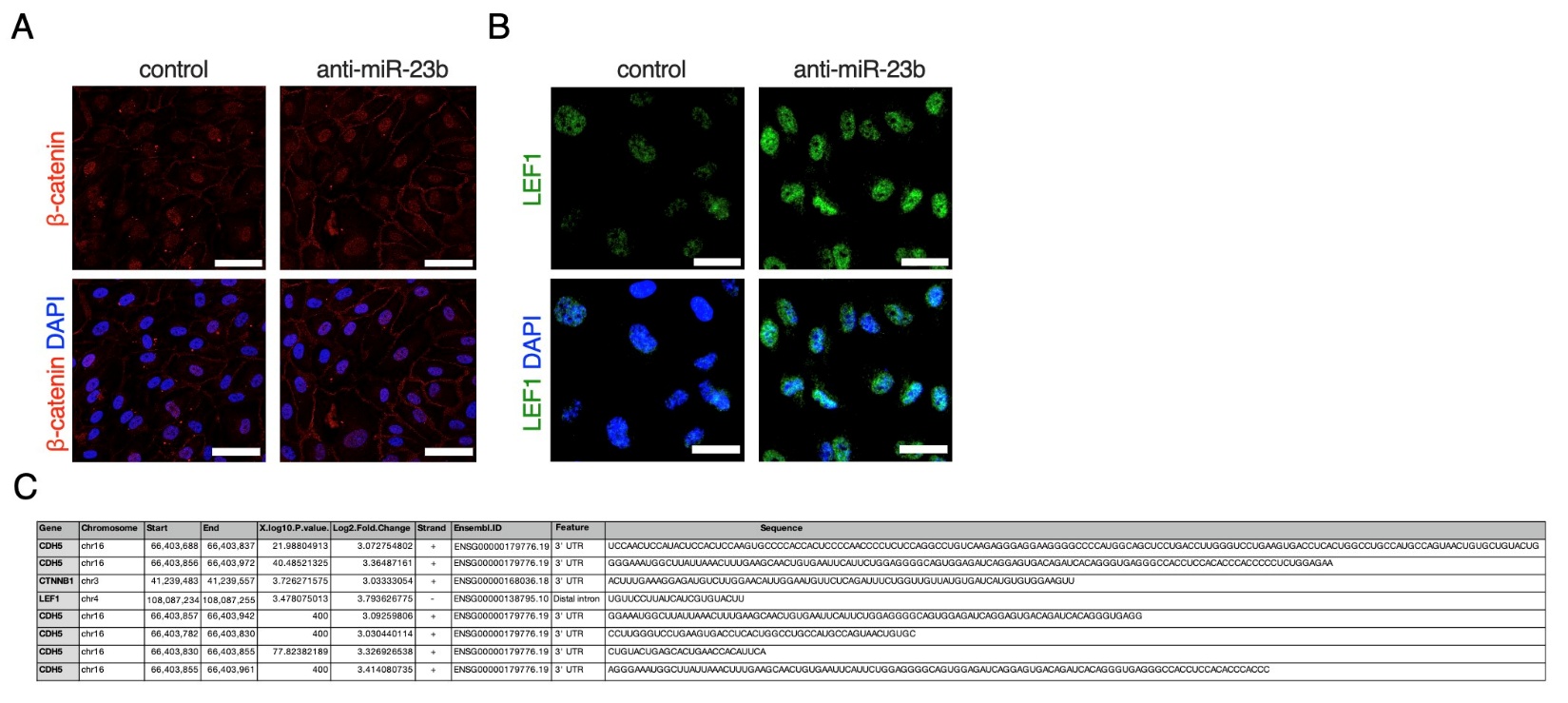

**Figure S4. HBMEC β-catenin and LEF-1 expression and eCLIP detail for miR-23b binding to CDH5, CTNNB1 and LEF-1 mRNA.** **A.** Magnification of representative IF images showing β-catenin and LEF-1 expression in HBMECs. Magnified image of β-catenin and LEF-1, scale = 50µm and 30µm.

**B.** miR-23b eCLIP-seq details delated to CDH5, CTNNB1 and LEF-1, including p-values, log2 fold change of miR-23b binding compared to control, strand bound, feature (3’UTR, distal intron) and sequence of immunopurified mRNAs bound to miR-23b.

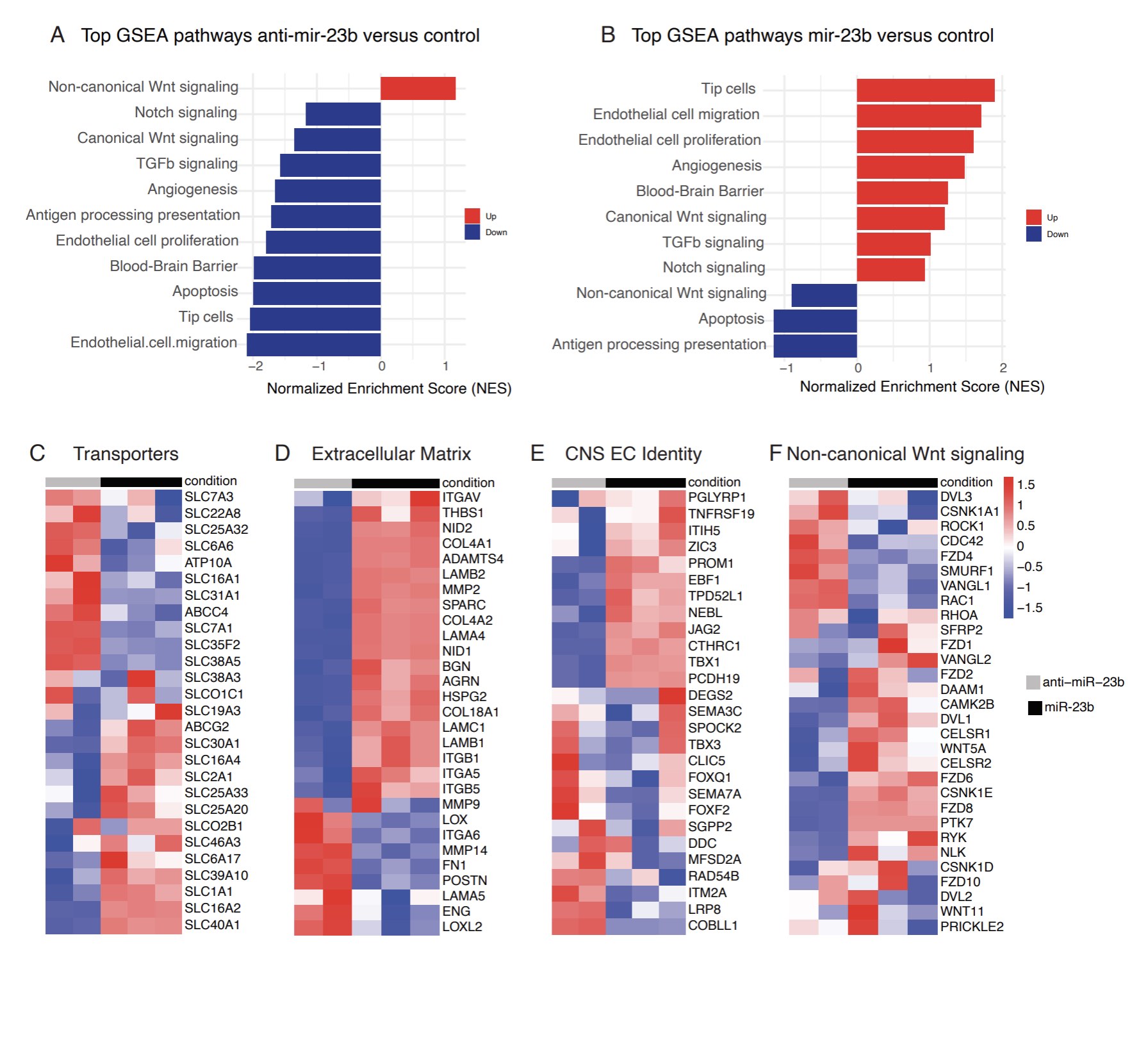

**Figure S5.1. anti-miR-23b and miR-23b affect BBB-associated pathways and CNS identity gene programs in HBMECs.** **A.** Gene Set Enrichment Analysis (GSEA) of enriched pathways comparing control miR and anti-miR-23b overexpressing HBMECs. Red bars indicate pathways upregulated in anti-miR-23b and blue bars indicate pathways enriched in control cells. **B.** Gene Set Enrichment Analysis (GSEA) of enriched pathways comparing miR-23b and control miR HBMECs. Red bars indicate pathways upregulated in miR-23b cells, and blue bars indicate pathways enriched in control miR cells. **C-F.** Heatmaps depicting differentially expressed genes associated with **C.** Transporters, **D.** Extracellular Matrix, **E.** CNS EC Identity, and **F.** Non-canonical Wnt signaling, between miR-23b and anti-miR-23b treated HBMECs, with the one outlier anti-miR-23b sample removed. Scale bar represents log(Z-score), where red indicates a high Z-score and blue indicates a low Z-score.

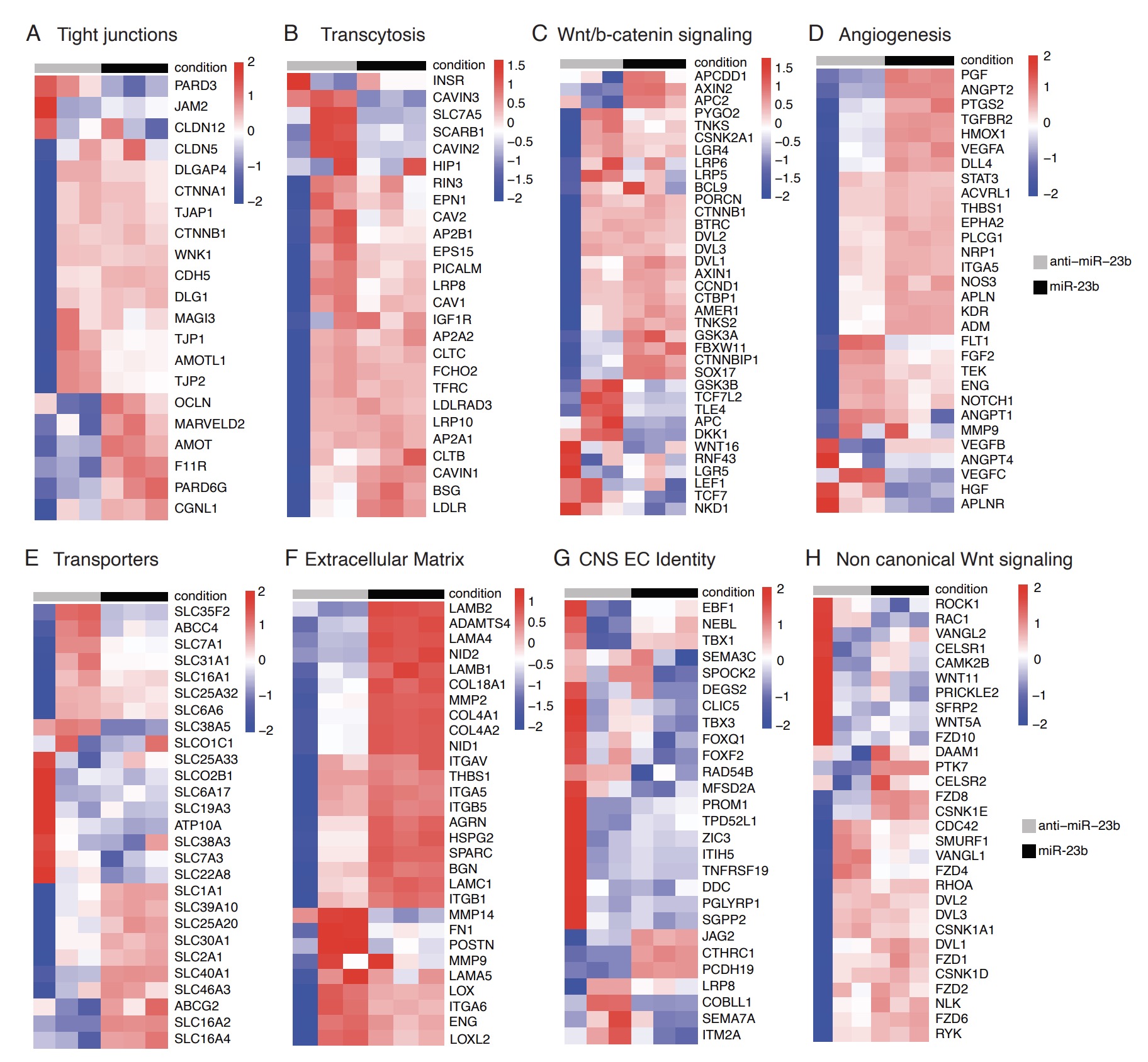

**Figure S5.2. anti-miR-23b and miR-23b affects BBB-associated pathways and gene programs in HBMECs. A-H.** Heatmaps depicting differentially expressed genes (including the outlier anti-miR-23b sample) associated with **A.** Tight Junction, **B**. Transcytosis, **C.** Wnt/β-catenin signaling, **D.** Angiogenesis, **E.** Transporters, **F.** Extracellular Matrix, **G.** CNS EC Identity, and **H.** Non-canonical Wnt signaling, between miR-23b and anti-miR-23b overexpressing HBMECs. Scale bar represents log(Z-score), where red indicates a high Z-score and blue indicates a low Z-score.

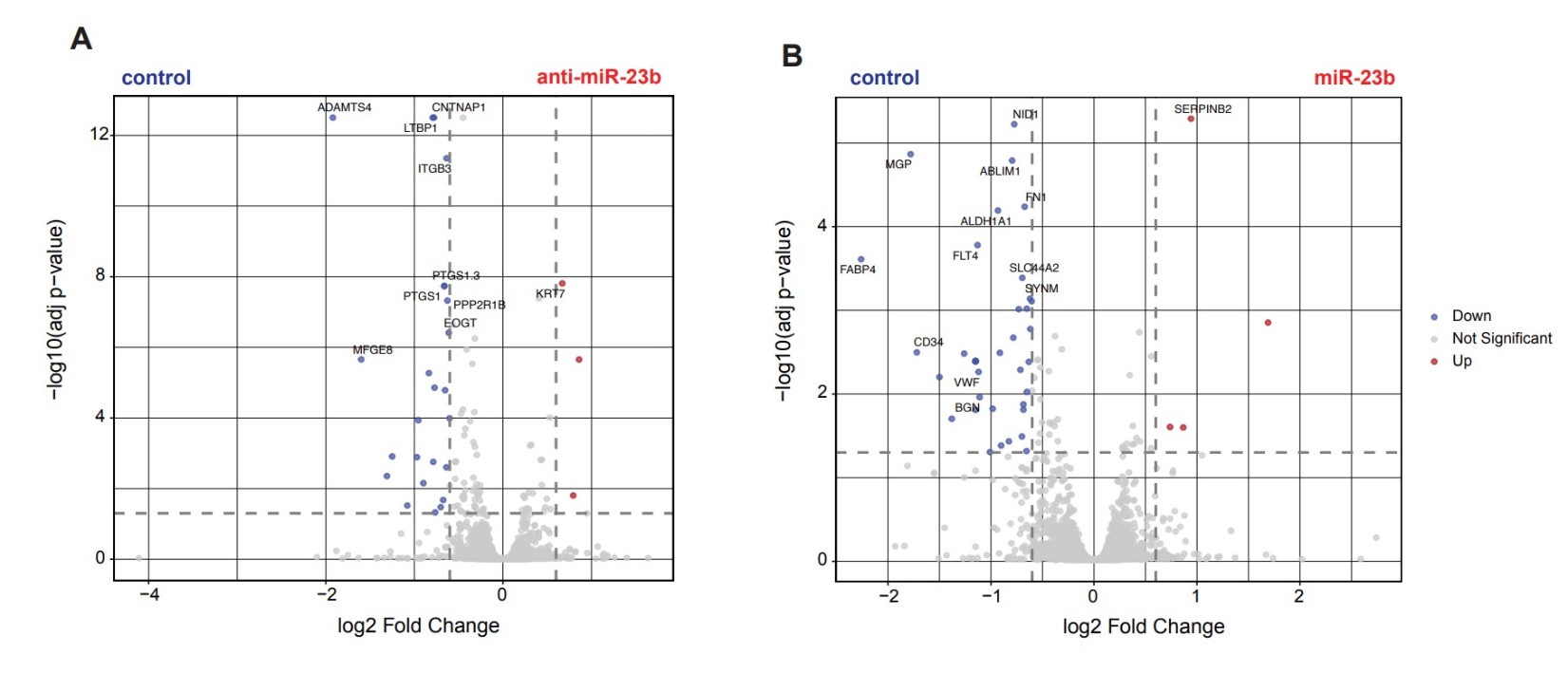

**Figure S5.3. anti-miR-23b and miR-23b modulation alters BBB-specific protein expression in HBMECs.** **A-B.** Volcano plot highlighting differentially expressed BBB-specific proteins identified by proteomic analysis between anti-miR-23b vs control and miR-23b vs control samples. All listed proteins are statistically significant (P~adj~ < 0.05; |log2FC| > 0.6).

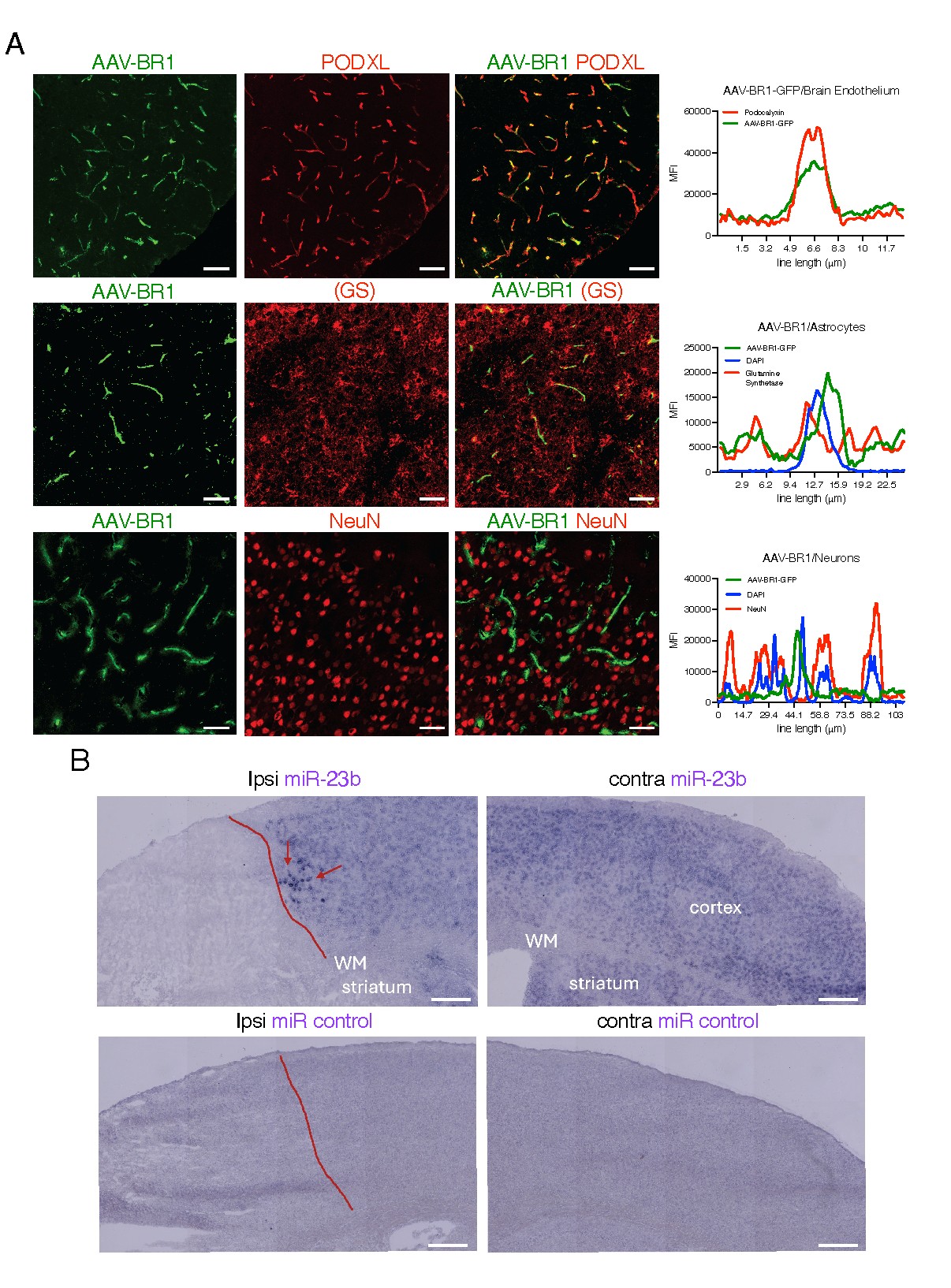

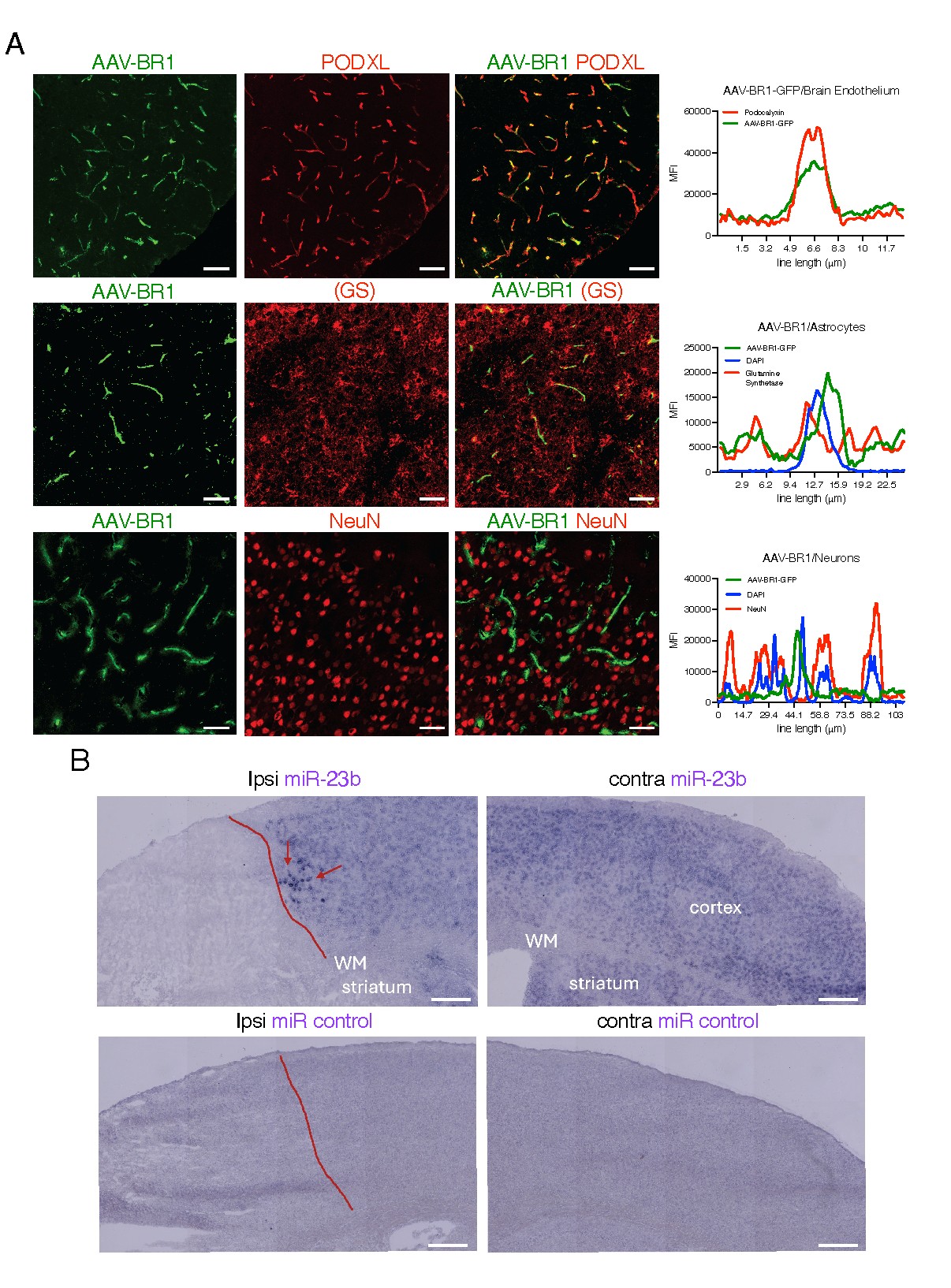

**Figure S6. AAV-BR1 expression in brain endothelial cells, in astrocytes and in neurons.**

**A.** Mice were transduced with AAV-BR1 control virus. A representative image of AAV-BR1-control-GFP (green) virus showing expression in brain capillaries co-stained with podocalyxin/PODXL (red, vessel marker), three weeks post-infection (top panel). Minimal (< 2%) AAV-BR1 expression was detected in astrocytes (GS positive) or neurons (NeuN positive). Right panels show the level of overlap between AAV-BR1 and PODXL, GS and NeuN. Scale = 100µm and 50µm (GS and NeuN). **B.** miR control and miR-23b in situ Hybridization was performed on t-MCAO tissue. Increased miR-23b expression is shown at the stroke region (red arrows), white matter (WM). Scale = 100µm.
